# A myosin filament cage drives multiple aspects of mitosis unique to cells dividing within a polarized epithelium

**DOI:** 10.64898/2026.10.25.761724

**Authors:** Sulaiman Yousafzai, Kirsten Remmert, Anjelika Gasilina, Grace Perry, Lelia Boley, Chris Alexander, Antonio Pedrosa, Dillon Schrock, Yang-In Yim, Xufeng Wu, John A. Hammer

## Abstract

Here we used mouse intestinal organoids expressing endogenously tagged myosin to clarify its roles in those aspects of mitosis unique to division within a polarized epithelium. Super-resolution imaging of rounded metaphase cells revealed a never-before-seen myosin filament cage that only forms in this context, exhibits apical/basal polarity, maintains roundness by resisting surrounding tissue pressure, and repositions laterally to form a contractile ring predetermined to drive planar division and asymmetric cleavage furrow progression. Attenuating cage function inhibited interkinetic nuclear migration and metaphase cell rounding, causing the division plane to shift from planar to orthogonal and resulting in organoids possessing multiple lumens. Following the apical domain marker ZO-1 showed that the new lumens originate at the interface between stacked daughter cells created by orthogonal divisions. We conclude that cells dividing within polarized epithelia create a myosin filament cage designed specifically to drive those aspects of cell division unique to their context.

## INTRODUCTION

Type 2 myosins are the major actin actin-based contractile motor protein in all non-muscle cell types [1-3]. Consistently, they are essential for a wide range of fundamental cellular processes that include cell division [4, 5], cell adhesion [6, 7], cell migration [8, 9], and tissue morphogenesis [10-13]. Myosin 2 drives these processes by assembling into small bipolar filaments [14]. When these myosin filaments encounter actin filaments of opposing orientation, they cause them to slide past each other, resulting in contraction of the actin cytoskeleton (much as muscle myosin does within the sarcomere).

The large number of studies of cells dividing in isolation all point to two major roles for myosin 2 during this process. First, it plays a central role in driving metaphase cell rounding, a universal feature of dividing cells, by creating a contractile, isotopically-distributed cortical actomyosin 2 network [15-19]. This network, together with an increase in hydrostatic pressure due to water influx, drives the isotropic rounding of metaphase cells [20, 21]. The primary purpose of metaphase rounding is to provide sufficient space for mitotic spindle formation and the optimal geometry for spindle positioning, microtubule: kinetochore attachment, and chromosome segregation [5, 22]. Consistently, isolated cells placed under vertical confinement so as to prevent rounding exhibit pronounced defects in spindle formation and function [23-25].

The other major role for myosin 2 in dividing cells is to form the contractile ring and then power its subsequent constriction to drive cleavage furrow ingression [26]. This function, when coupled with abscission at the midbody, separates the daughter cells [27, 28]. Creation of the contractile ring is thought to involve a combination of de novo assembly at the midzone and a cortical flow of preexisting, peripheral actomyosin assembles that then rearrange at the midzone [16, 29-31]. The de novo assembly pathway is driven by the spindle microtubule-dependent activation of ECT2, a guanine nucleotide exchange factor for RhoA [32-34]. GTP-loaded RhoA then activates the formin-dependent assembly of linear actin filaments and the Rho-kinase dependent assembly of myosin 2 filaments at the midzone during anaphase [35, 36]. The division plane in cells dividing in isolation is typically perpendicular to the cell’s long axis just prior to mitosis (Hertwig’s rule). Finally, the contractile ring in cells dividing in isolation closes in largely symmetric fashion.

Importantly, cells dividing within a polarized epithelium must undergo several myosin dependent processes that cells dividing in isolation do not do. First, the nucleus, which resides during interphase at the base of the cell, must be moved apically to the cell center in a process called interkinetic nuclear migration [37-39]. Second, the division plane must be positioned perpendicular to the epithelial plane so as to ensure a planar cell division, as such divisions maintain epithelial integrity by forcing the daughter cells to remain in the epithelial plane [40, 41]. Third, the contractile ring must close asymmetrically in basal to apical fashion so as to maintain barrier function throughout division and to allow in coordinated fashion the forming of new cell: cell contacts between the daughter cells [42, 43]. Finally, while cells dividing within a polarized epithelium share with cells dividing in isolation the need to round up during metaphase, the advantages of rounding play outsized roles for cells dividing within a polarized epithelium because, unlike cells dividing in isolation, cells within the epithelium must deal with compressive forces coming from the surrounding tissue [44]. Taken together, then, all of the major aspects of cell division for cells dividing within a polarized epithelium-nuclear positioning, metaphase rounding, setting the division axis, and cleavage furrow progression- are unique relative to cells dividing in isolation.

Here we sought a deeper understanding of myosin 2’s role in these unique aspects of cell division by combining its endogenous tagging with super resolution microscopy in the context of a 3D, stem cell-driven, polarized epithelial cell model. These efforts lead to the identification of a dynamic myosin filament cage that only forms in cells dividing within a polarized epithelium, exhibits apical/basal polarity, and appears specifically designed to drive multiple aspects of mitosis unique to cells dividing in this context, including resistance to compressive forces, a planar axis of division, and asymmetric cleavage furrow progression. Attenuating cage function causes the division plane to shift from planar to orthogonal and results in organoids possessing multiple lumens. These and other results shed important new light on the mechanism of cell division within polarized epithelia and the consequences of its disruption.

## RESULTS

### The interphase localizations of myosin 2A and myosin 2C, the only isoforms expressed in enterocytes, suggest largely distinct functions

We first sought to determine which myosin 2 isoforms are expressed in intestinal organoids and where they localize during interphase. Mice possess three myosin 2 genes-Myh9, Myh10, and Myh14-that encode the heavy chains for myosins 2A, 2B and 2C, respectively (referred to below as M2A, M2B and M2C). These three isoforms differ as regards their kinetic and mechanical properties, often exhibit distinct localizations, and have both overlapping and unique functions depending on the cellular context [45]. To determine which of these M2 isoforms are expressed in intestinal epithelial cells, we imaged sections of small intestine prepared from three mouse lines in which GFP had been knocked into the endogenous M2 heavy chain locus by homologous recombination. We refer to these three lines throughout the text as GFP-M2A knockin (KI), GFP-M2B KI, and GFP-M2C KI mice. All three lines are maintained as homozygous KIs, and GFP is present at the N-terminus of M2A and M2B, and at the C-terminus of M2C. Consistent with immunostaining in previous studies [10], tissue sections reveal bright signals for M2A and M2C, but not for M2B (Fig. 1, Panels A1-A3). Similarly, organoids exhibit bright signals for M2A and M2C, but not for M2B (Fig. 1, Panels B1-B3). Consistent with these images, Western blots of small intestine extracts probed with isoform specific antibodies show clear signals for M2A and M2C but not for M2B (Fig. 1, Panel C). While we cannot exclude some role(s) for M2B, all subsequent efforts focused only on M2A and M2C.

**Figure 1.**
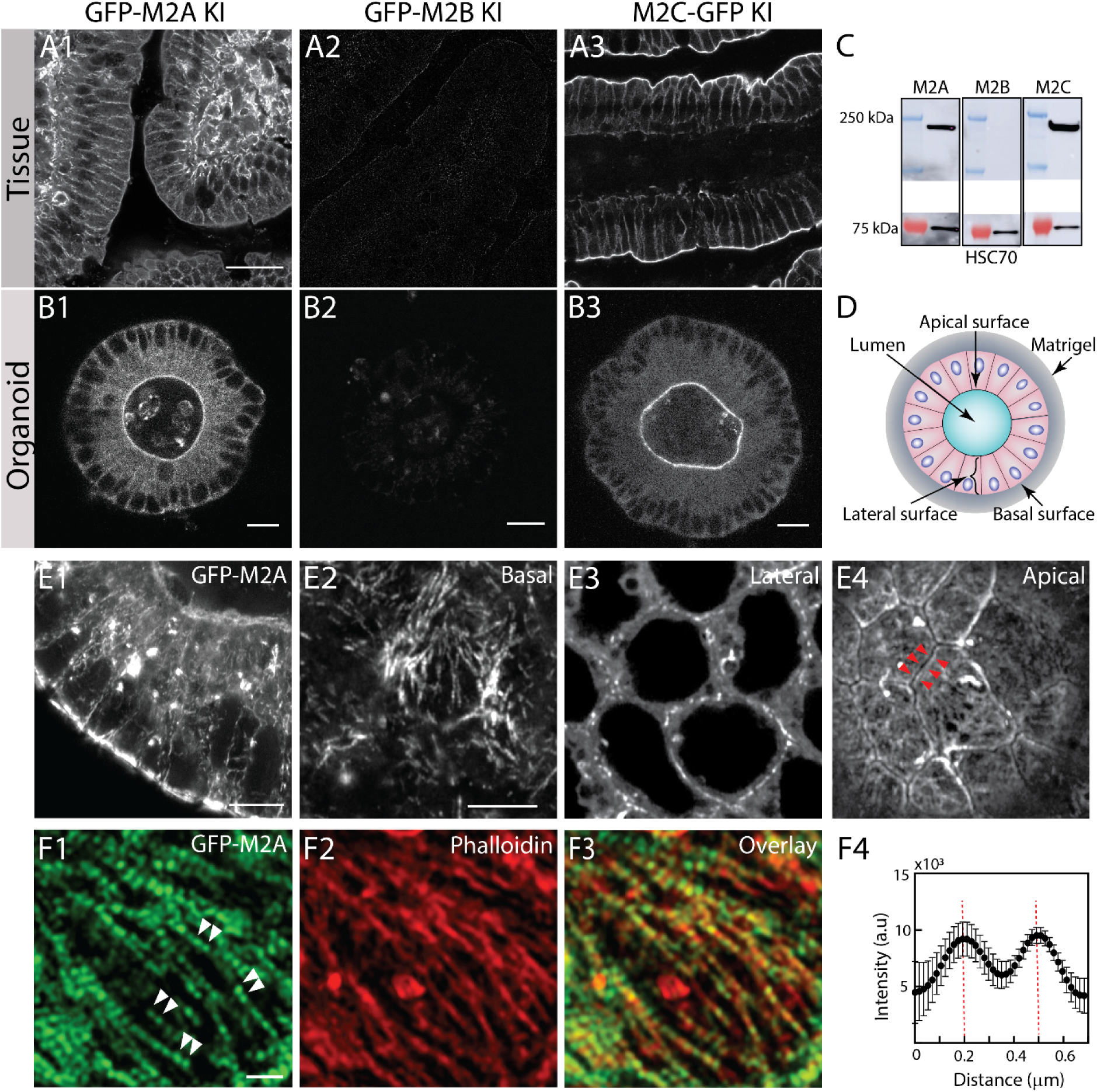
The interphase localization of endogenously tagged M2A is consistent with it supporting cell: cell and cell: ECM adhesion. (A, B) Representative confocal images of small intestinal tissue (A1-A3) and organoids (B1-B3) isolated from GFP-M2A (A1, B1), GFP-M2B (A2, B2), and M2C-GFP (A3, B3) KI mice. (C) Western blots of whole-organoid lysates probed with isoform-specific antibodies against the ∼200 kDa heavy chains of M2A, M2B and M2C (top), and with an antibody against Heat Shock Protein 70 (HSC70) (bottom) as the loading control. (D) Schematic of an intestinal organoid embedded within Matrigel (gray) depicting its single lumen (blue) surrounded by a single layer of enterocytes (pink). Arrows mark the apical, lateral and basal surfaces of the enterocytes. (E1) Representative, Z-projected AiryScan image of a subset of equatorial planes acquired from a GFP-M2A KI organoid quadrant using 0.2 µm steps over a ∼20 µm depth. See also Movie 1 for a representative basal surface to apical surface Z-stack. (E2) Representative AirSycan confocal image (single plane) of the basal surface. (E3) Representative AiryScan confocal image (5 µm Z-projection) of the lateral surface. (E4) Representative AiryScan confocal image (single plane) of the apical surface (the red arrowheads mark the moderate enrichment of GFP-M2A along bicellular junctions). With regard to GFP-M2A along the lateral surface, see also Movie 2 for a Z-Stack and Movie 3 for a 3D rendering. (F1) Representative AiryScan confocal image of the GFP-M2A signal (single plane) across the cell’s basal surface. The double white arrowheads mark the two ends of individual M2A bipolar filaments, which exhibit ∼300-nm spacing due to M2A being tagged with GFP at its N-terminus (see refs [1, 46] and Panel F4). (F2) Signal for Alexa 488-labeled Phalloidin in this same confocal section. (F3) Overlaid image of these M2A-decorated basal stress fibers. (F4) Quantification of line-scan intensities along the long axis of the GFP-M2A bipolar filaments. Scale bars: (A1) 30 µm; (B1-B3) 10 µm; (E1) 5 µm; (E2-E4) 5 µm; (F1) 1 µm.

We used optical sectioning of fixed organoids (Fig. 1, Panel D) made from KI mice to determine the steady state localizations of endogenously tagged M2A and M2C in interphase cells. Maximum projection images of GFP-M2A KI organoids show that this isoform is enriched primarily along the basal and lateral surfaces of the epithelium (Fig. 1, Panels E1). More insightful are Z-Stack movies and representative confocal sections (Fig. 1, Panels E2-E4; see also Movie 1). Sections at the cell base, the site of cell: ECM adhesion, show robust localization of M2A in the form of long stands that traverse most of the ventral surface (Fig. 1, Panel E2). Higher magnification images show that these strands are composed of individual bipolar filaments of M2A in linear register, as evidenced by repeating doublets of GFP spaced ∼300 nm apart (the signature for N-terminally tagged M2A; [1, 46, 47](Fig. 1, Panels F1 and F4). As expected, staining for F-actin (Fig. 1, Panels F2 and F3) shows that these strands correspond to stress fibers. The strain that these M2A-rich stress fibers place on these adhesions likely serves to keep enterocytes tightly bound to the Matrigel by keeping their integrins in the open, active conformation [48]. Sections along the lateral surface of enterocytes, the site of cell: cell adhesion, show strong albeit discontinuous M2A localization (Fig. 1, Panels E1 and E3). This is best shown in Z-stacks (Movie 2) and 3D renderings (Movie 3) that span the lateral surfaces. This localization is consistent with the known roles that type 2 myosins play in linking adherens junctions, which drive cadherin-based cell: cell adhesion, to the actin cytoskeleton. Finally, sections at the cell apex show that M2A is slightly enriched where the apical membranes of adjacent enterocytes abut (Fig. 1, Panel E4; see red arrowheads). Together, these observations are consistent with M2A playing major roles during interphase in supporting cell: ECM and cell: cell adhesion, and possibly some role in supporting tight junction organization/function.

Consistent with the low magnification images in Figure 1, as well with images in a recent study [49], maximum projection images of GFP-M2C KI organoids show that M2C localizes almost exclusively at apical junctions (Fig. 2, Panel A1). Moreover, staining for ZO-1 reveals a dramatic correspondence between this tight junction marker and the signal for M2C (Fig. 2, Panels A2-A4). High magnification, maximum projection images show that M2C is present in the form of dots lining the ZO-1-marked apical perimeter of enterocytes (Fig. 2, Panels B1, B2 and B4). These M2C dots are spaced roughly evenly across all bicellular junctions (see the bracket in Panel B2), and line scans across these dots reveal a periodicity of ∼400 nm (Fig. 2, Panel B3), consistent with the spacing of C-terminally tagged M2C filaments in linear register [1, 50]. Finally, the M2C dots appear brighter at tricellular junctions (four are boxed in Fig. 2, Panel B4). This later point is further highlighted in high mag insets (Fig. 2, Panel 5). Together, these observations suggest that M2C’s primary role during interphase is probably to support barrier function by controlling tight junction organization and/or function.

**Figure 2.**
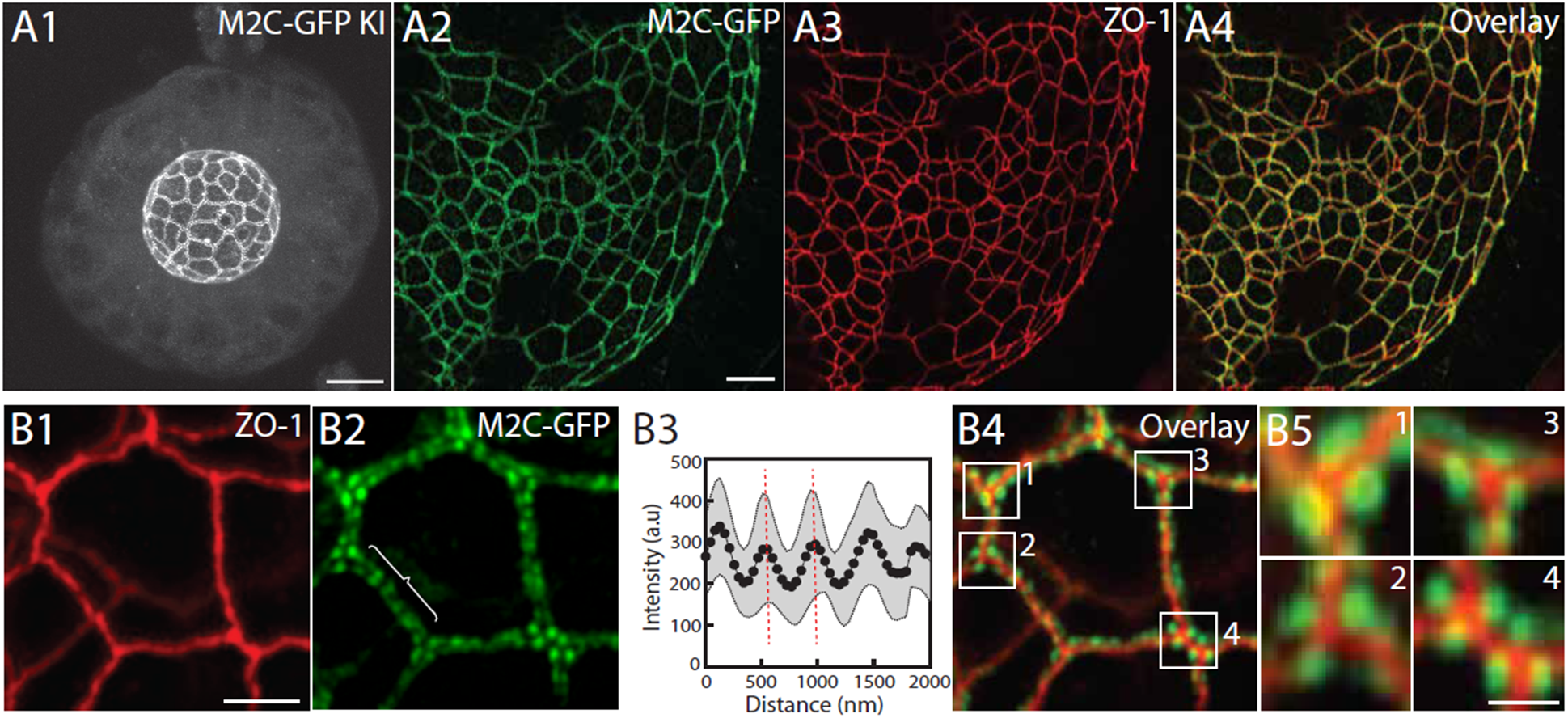
The localization of endogenously tagged M2C is consistent with it regulating tight junction structure and/or function during interphase. (A1) Representative, maximum-intensity projected Airyscan confocal image of an entire M2C-GFP KI organoid. (A2) Representative, maximum intensity projected AiryScan image of the apical surface of a M2C-GFP KI organoid. (A3) The corresponding signal for the apical domain tight junction marker ZO-1 (by immunostaining). (A4) Overlay of the signals in A2 and A3. (B1) Magnified image of ZO-1 staining. (B2) Corresponding magnified image of M2C-GFP. (B3) Quantification of line-scan intensities for the M2C signals along bicellular junctions. The ∼400 nm spacing is due to M2C being tagged with GFP at its C-terminus (so the spacing reports the center of adjacent bipolar filaments). (B4) Overlay of the M2C-GFP and ZO-1 signals in Panels B1 and B2, with four tricellular junctions boxed. (B5) Higher magnification images of the four boxed regions in Panel B4. Scale bars: (A1) 10 μm; (A2) 5 μm; (B1) 2 μm.

### M2A accumulates in structures central to driving cell division in polarized epithelia

We next used diffraction-limited, time lapse imaging of dividing cells in GFP-M2A and GFP-M2C KI organoids transduced with Lenti-RFP-Histone 2B to look for the presence of these two isoforms in key actomyosin-dependent mitotic structures. With regard to M2A, its first site of accumulation is under the nucleus during interkinetic nuclear migration (INM) (Fig. 3; compare Panels A1/B1 (“Interphase”) to Panels A2/B2 (“INM”); the yellow arrowheads in Panel B2 mark the accumulation of GFP-M2A; see also Movie 4). The contractile force generated at this site creates not only a pushing force that moves the nucleus in the apical direction but also a pulling force that results in a pronounced invagination beneath the dividing cell (compare the dashed white lines marking the outer perimeter of the organoid in Panels B1 and B2). While this invagination is accompanied by significant shrinkage of the cell’s base, and may involve some amount of focal adhesion disassembly [51], the dividing cell remains firmly attached to the ECM throughout mitosis (see also below). As INM approaches completion, M2A assumes a uniform cortical localization to drive metaphase cell rounding (Fig. 3, Panels A3 and B3 (“Metaphase Rounding”); the yellow arrowheads in Panel B3 mark the even cortical accumulation of GFP-M2A; see also Movie 4). Finally, M2A accumulates between the separating chromosomes at anaphase to form the contractile ring (Fig. 3, Panels A4 and B4 (“Contractile Ring”); the yellow arrowheads in Panel B4 mark the accumulation of GFP-M2A in the forming ring; see also Movie 4). In contrast to these results, time lapse images of dividing cells in GFP-M2C KI organoids revealed little to no accumulation of M2C in any of these mitotic structures (see Movie 5). Given that, and given that M2C KO organoids do no exhibit an obvious defect in cell division (see below), all subsequent efforts to characterize the organization and dynamics of myosin filaments during mitosis using super resolution imaging focused on M2A.

**Figure 3.**
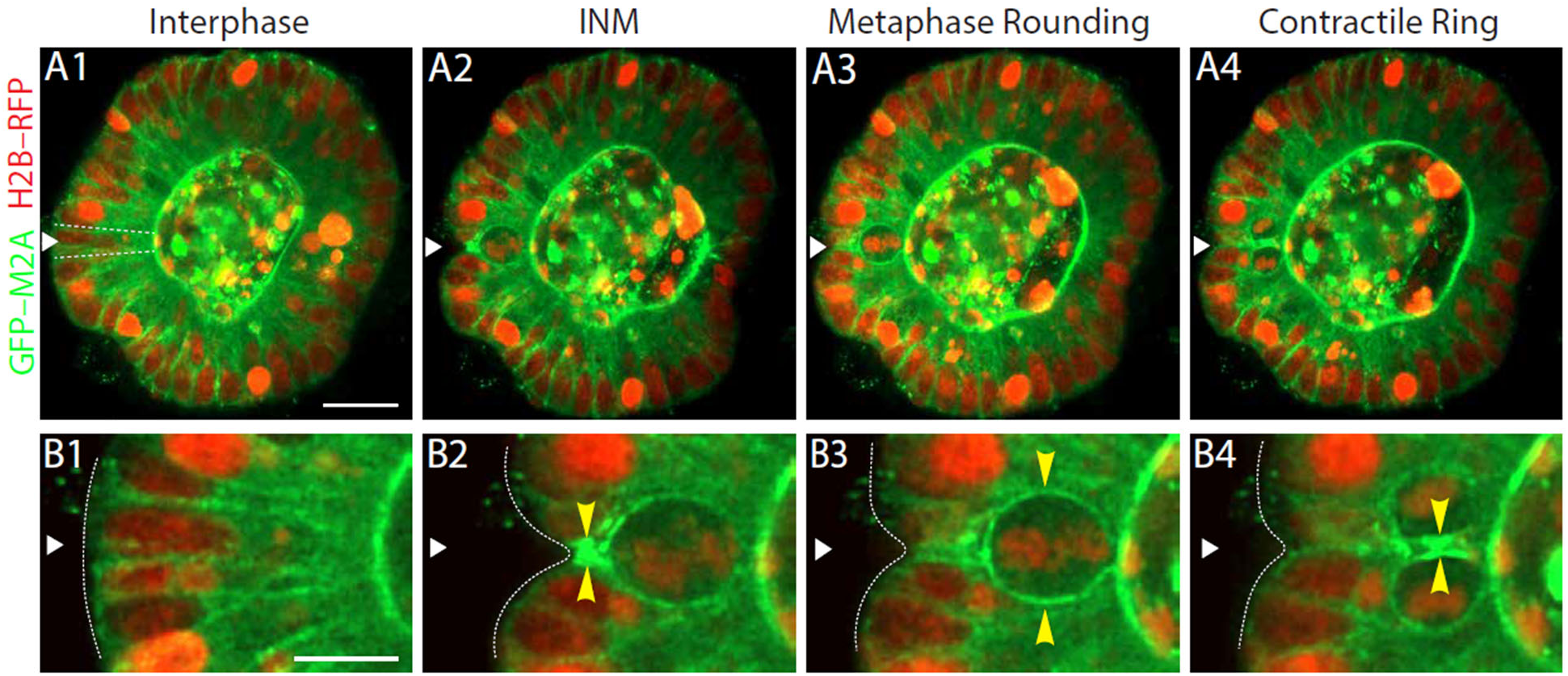
M2A accumulates in structures central to driving cell division within polarized epithelia. (A, B) Snapshots from a representative, diffraction-limited, time lapse movie of a dividing cell within a GFP-M2A KI organoid expressing H2B-RFP to mark DNA. (A1-A4) The dividing cell (marked by a white arrowhead in all four panels and outlined by a dashed white line in Panel A1) during interphase (A1), during INM (A2), during metaphase rounding (A3), and during contractile ring formation (A4). (B1-B4) Corresponding magnified images of this dividing cell. The yellow arrowheads in B2-B4 point to GFP-M2A accumulations at each step, and the white dashed line in B1-B4 outlines the organoid’s basal surface. See also Movie 4. Scale bars: (A1) 20 µm; (B1) 10 µm.

### Metaphase cells possess an evenly-spaced cortical cage of M2A filaments that exhibits apical basal polarity

Previous efforts to image M2 distribution in the cortex of rounded metaphase cells dividing in isolation (reviewed in [5, 15]), as well the diffraction-limited, time lapse imaging of M2A in cells dividing within an organoid described above, are all consistent with the myosin being distributed in isotropic fashion and in the form of an actomyosin meshwork. Super resolution, Airyscan imaging of metaphase cells in fixed organoids revealed, on the other hand, M2A filaments arranged in strings running apically to basally (Fig. 4, Panels A1 and A2; A1 is a projection of several confocal sections; A2 is a single section from this stack; the red doublet arrowheads mark individual M2A bipolar filaments exhibiting the expected ∼300 nm periodicity). 3D renderings show that these M2A filament strings line the cortex of rounded metaphase cells (Movies 6 and 7), and lines scans across this M2A filament “cage” reveal a relatively uniform spacing of ∼1.25 µm between the adjacent filament strings (Fig. 4, Panels B1 and B2). Moreover, measurements of the axial orientation of individual M2A filaments within each string, determined by measuring the variance in the angles of lines drawn through the long axis of each bipolar filament, confirm what is clear to the eye, i.e. that filaments within each string are highly aligned relative to their long axis (Fig. 4, Panel C). Unlike fixed samples, individual filaments are not quite resolvable in dynamic, 3D Airyscan imaging of live metaphase GFP-M2A KI cells, presumably because of motion-dependent blurring occurring during image acquisition. That said, such dynamic imaging shows clearly the string-like organization of M2A filaments in the cortex of live metaphase cells (Movie 8). Finally, phalloidin staining shows that, as anticipated, strands of F-actin colocalize with the M2A filament strings comprising the cage (Fig. 4, Panels G1-G3).

**Figure 4.**
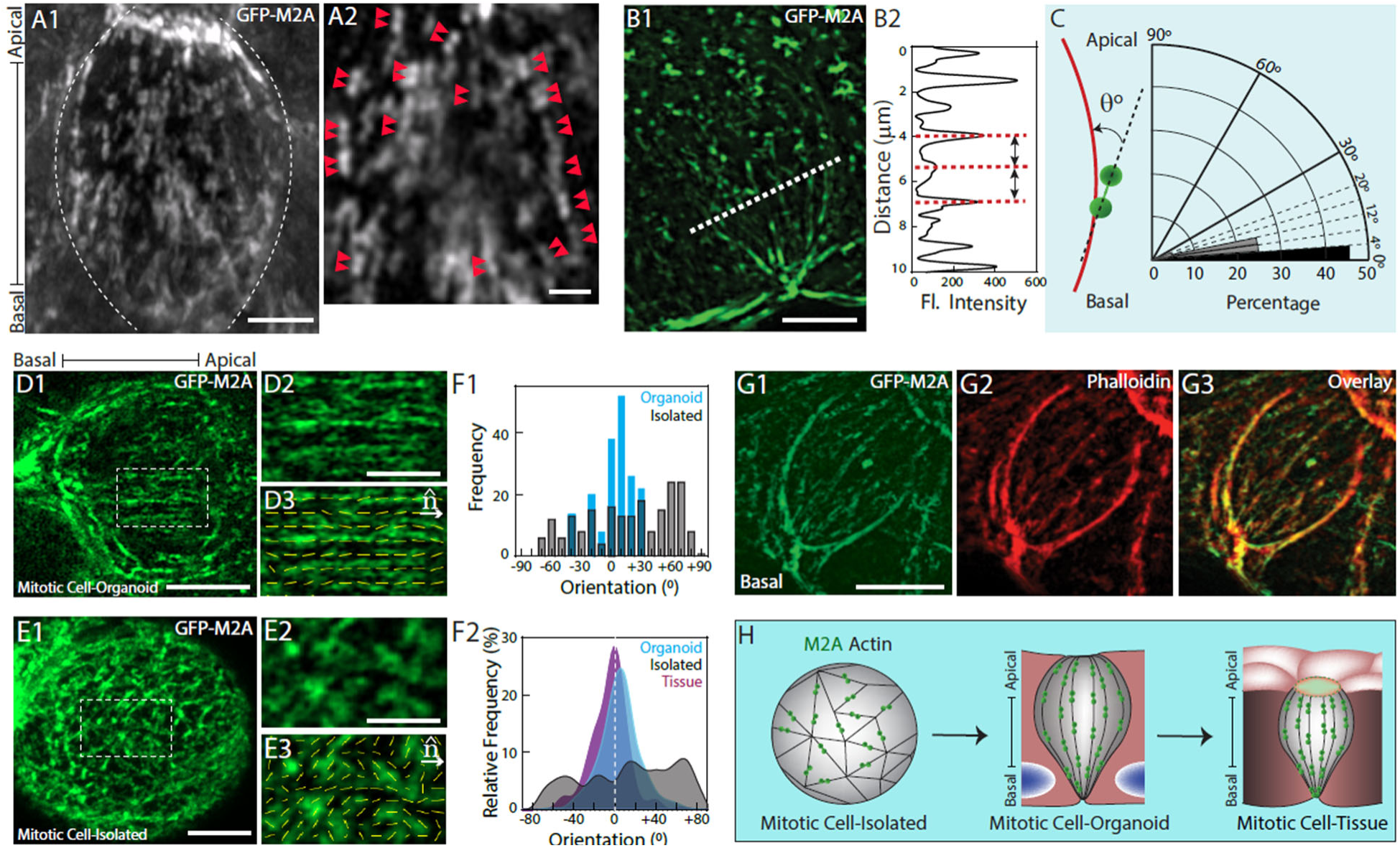
Metaphase cells within a polarized epithelium possess a cortical M2A filament cage composed of filament strings running apically to basally. (A1) Representative, maximum intensity, projected AiryScan image (4 µm Z-projection) of a rounded metaphase cell (outlined by the dashed white line) within a GFP-M2A KI organoid. (A2) A single section from this Z-stack (the double red arrowheads mark the two ends of individual M2A bipolar filaments). See also Movies 6 and 7 for 3D renderings of this metaphase M2A filament cage. (B1) Lower resolution, projected image of a M2A filament cage. (B2) Line-scan for GFP-M2A fluorescence across the portion of the cage in B1 that is most in focus (see the dashed line in B1). (C) Measurements of the axial orientation of individual M2A bipolar filaments within 10 filament strings. (D-F) The orientations of M2A filament long axes show that the metaphase M2A filament cage only forms when enterocytes divide within a polarized epithelium. (D1) Representative metaphase cell dividing within a GFP-M2A KI organoid (the image is oriented such that the cell’s apical/basal axis is horizontal). (D2) An enlargement of the boxed region in D1, which is most in focus. (D3) The long axis of all the M2A bipolar filaments within this boxed region are marked with yellow lines. (E1-E3) Similar to D1-D3 except that this enterocyte is dividing in isolation (and, as a consequence, lacks apical/basal polarity). (F1) Histograms for the orientations of all the marked filaments in D3 (blue) and E3 (gray) relative to the direction vector ***n̂*** (an orientation of 0° is parallel to this direction vector, while values of -90° and +90° are perpendicular and in opposite orientations to vector ***n̂***). (F2) The cumulative GFP-M2A filament angular orientations for 10 metaphase cells within an organoid (blue), 10 metaphase cells in isolation (gray), and 10 metaphase cells within tissue (maroon) (see Figure S1 for a representative example of the metaphase myosin filament cage within tissue). (G1) Maximum i ntensity projection of sixteen AiryScan confocal sections spanning an 8µm depth of a representative metaphase cell within a GFP-M2A KI organoid. (G2) The same sample stained with Phalloidin. (G3) The overlaid image. (H) A cartoon depicting the metaphase organization of actin and M2A filaments in cells dividing in isolation, within an organoid, and within tissue. Scale bars: (A1) 2 µm; (A2) 1 µm; (B1) 5 µm; (D1, E1) 5 µm; (D2, E2) 2 µm; (G1) 5 µm.

### The metaphase M2A filament cage only forms when cells divide within a polarized epithelium

To our knowledge, the metaphase myosin filament cage shown above has not been described previously in dividing epithelial cells. This could be due in significant part to our use of endogenously tagged GFP-M2A rather than transfected, over-expressed GFP-tagged M2A. Endogenous tagging is not the whole story, however, as the cage is only evident in metaphase GFP-M2A KI cells using super resolution microscopy (compare the diffraction limited image in Fig. 3, Panel B3, to the AiryScan super resolution images in Fig. 4). Given that the majority of previous efforts to image M2 in dividing cells were done using isolated cells rather than cells within an epithelial layer, we thought is also possible that the cage only forms when cells divide within an epithelial layer. To test for this, we compared the organization of M2A filaments in GFP-M2A KI cells dividing within an organoid epithelium (Fig. 4, Panel D1) versus dividing in isolation (Fig. 4, Panel E1; Panels D1 and E1 are representative, maximum-intensity projections of several consecutive AiryScan slices; the boxed regions within both, which are enlarged in Panels D2 and E2, respectively, correspond to a region of the metaphase cortex that is most in focus). While most of the individual M2A bipolar filaments within these images are not resolvable as ∼300 nm doublets, their long axis is still clear, allowing their orientation relative to axis of the image to be determined (Fig. 4, Panels D3 and E3; the yellow lines in these orientation fields denote the long axis of all individual filaments within the field). The blue bars in Fig. 4 Panel F1 (“Organoid”) show the distributions of the filament long-axis orientations relative to the basal/apical axis of this dividing enterocyte for all of the filaments in Panel D3, while the gray bars show the same for all of the filaments in Panel E3 relative to the horizontal axis of the image (note that the angles must fall between −90° and +90°). Consistent with visual inspection of Panel D3, the long axis orientations of most filaments in this organoid sample fall within a fairly narrow distribution (∼ ± 30°) that exhibits a major peak centered near 0°. In other words, the long axis orientations of most M2A filaments in the cortex of this metaphase cell align closely with the cell’s apical/basal axis. Conversely, and consistent with visual inspection of Panel E3, the long axis orientations of filaments in this isolated cell sample are spread roughly evenly across a broad distribution relative to the horizontal axis of the image. In other words, the long axes of most M2A filaments in the cortex of this isolated metaphase cell exhibit very little uniformity as regards their orientation. These results were confirmed by quantitating the angular orientations of M2A filaments in ten metaphase cells for each condition (Fig. 4, Panel F2; Blue, metaphase cells in an organoid; Gray, isolated metaphase cells), as well as in ten metaphase cells present within tissue sections from the GFP-M2A KI mouse (Fig. 4, Panel F2; Magenta; see Fig. S1). Together, these results argue that the metaphase M2A filament cage only forms when cells divide within a polarized epithelium (see the cartoon on Fig. 4, Panel H). This finding suggests that cage formation requires the polarity cues that cells are provided only when they divide in that context. For example, the polarity cues may dictate the localization of the actin assembly machinery (most likely one of more formins) required to create the cage. Even more importantly, this finding suggests that the M2A filament cage may possess design features specifically suited to drive those aspects of cell division unique to cells dividing in a polarized epithelium.

### The metaphase myosin filament cage serves to resist surrounding compressive forces

Metaphase cell rounding in isolated cells is the consequence of two opposing forces-an isotropic swelling force driven by hydrostatic pressure and an isotopic contractile force driven by a meshwork of actomyosin 2 distributed evenly throughout cell’s cortex [20]. Consistently, global inhibition of myosin function in isolated metaphase cells results in their isotropic expansion [20]. For cells dividing within a tissue like the organoids studied here, external compressive forces arising from surrounding cells and tissue represent a third force effecting metaphase cell shape. Given this, one would predict that localized inhibition of the myosin cage would cause the cell to invaginate at that site, to exhibit a compensatory expansion elsewhere, and to lose roundness if these changes are not equal. To test for this, we performed UV laser ablation on one side of GFP-M2A-expressing metaphase cells along their apical/basal axis (Fig. 5). Of the thirteen ablations performed, the cells exhibited minimal morphological change in two cases and apparent rupture in two cases (Fig. 5, Panel A). Of the remaining nine cases, every cell exhibited a pronounced movement away from the ablation site i.e. none moved towards the ablation site (Fig. 5, Panel A; Panels B1-B4 show a representative example of the movement following ablation; the dashed white line in all four panels shows the cell’s shape just prior to ablation while the red dashed lines show that cell’s shape 40s and 80s after ablation; see also Movie 9). Consistent with the shape of the dashed red line at 80s, particle image velocimetry (PIV) analysis (Fig. 5, Panel C) showed that the cortical displacement on the ablated side of this cell was significantly larger than on its opposite, non-ablated side, indicating asymmetric cortical relaxation. Such differential surface motions should result in loss of roundness. To prove this, we measured the curvatures on both sides of the nine successfully ablated cells by fitting circles of radii R1 and R2 to the ablated and non-ablated surfaces, respectively, so as to approximate each surface as locally spherical (Fig. 5, Panel D1). Quantitative analysis showed that the ablated cells do indeed lose roundness (Fig. 5, Panel D2). Together, these findings argue that the metaphase myosin filament cage serves to promote near-isotropic rounding and mechanical stability in the context of a mechanically constrained epithelium.

**Figure 5.**
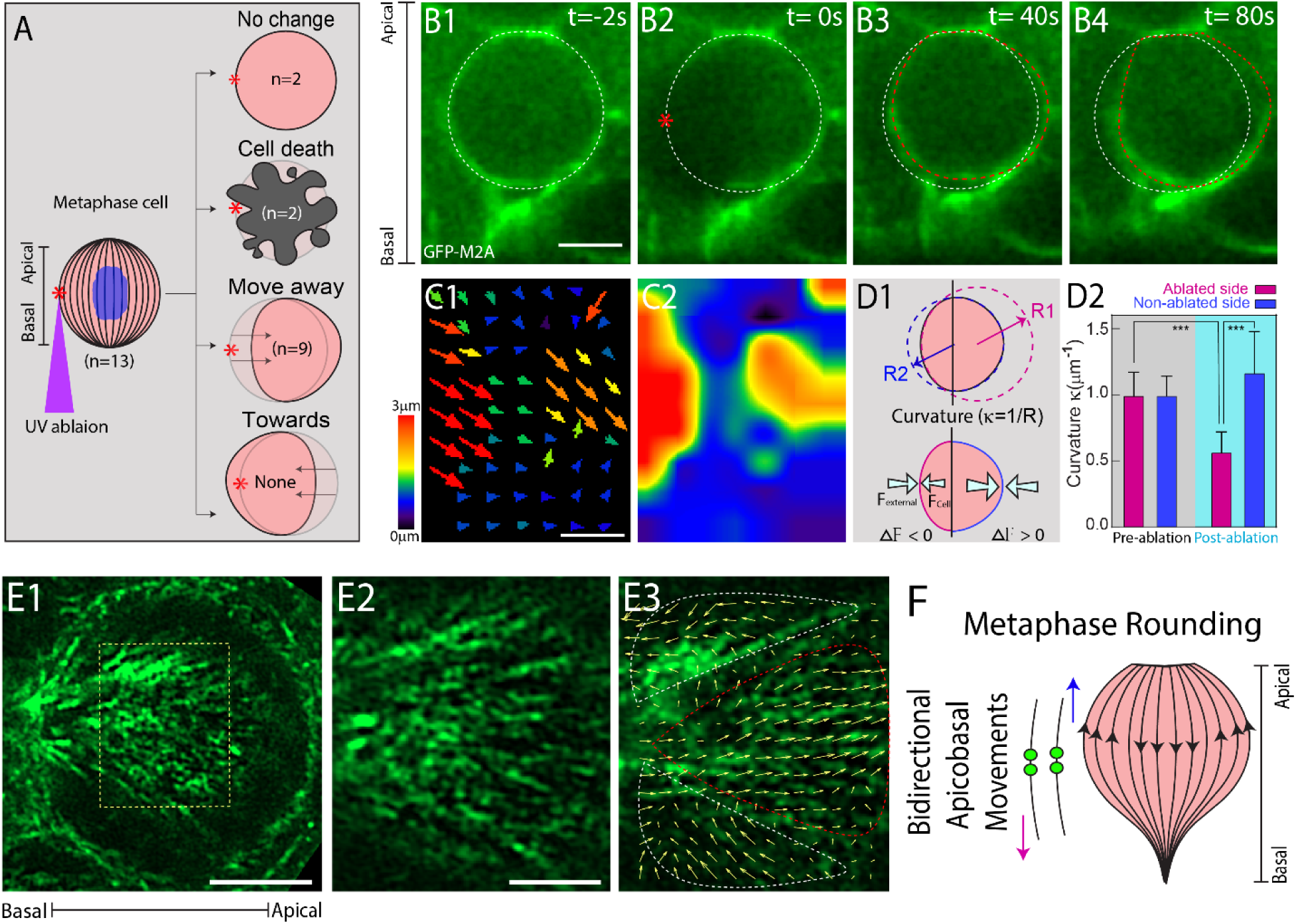
The cortical M2A filament cage supports metaphase rounding in the presence of external compressive forces through bidirectional apicobasal contractility. (A) Schematic of how the UV laser was used to ablate the lateral edge of the metaphase cell’s cortex (red asterisk) for 13 cells dividing within a GFP-M2A KI organoid. Post-ablation responses were categorized as no change (n = 2), cell death (n = 2), displacement away from the ablation site (n = 9), and movement toward the ablation site (n = 0). (B1-B4) Stills from a representative time lapse movie where the ablated cell moved away from the ablation site (the white dashed lines in all four panels note the outline of the rounded metaphase cell 2 seconds before ablation) (B1), while the dashed red lines in B2-B4 note the cell outline at the moment of ablation, 40 seconds post ablation, and 80 seconds post abolition, respectively (see also Movie 9). (C1) Particle Image Velocimetry (PIV) for the cell displaced in B1-B4 (the arrows show the total displacement over 80 seconds). (C2) Heatmap of the displacement magnitudes in C1. (D1) Schematic illustrating how fitting circles to the ablated side (R1 red) and to the non-ablated side (R2, blue) was used to quantitate changes in curvature (κ = 1/R) following ablation. (D2) The cumulative changes in curvature before and 80 seconds after ablation for the 9 cells that moved away from the ablation site (p < 0.001). (E) Groups of myosin filaments move simultaneously and bidirectionally along the metaphase cell’s apical/basal axis. (E1) Still from a representative time lapse Airyscan movie of the cortex of a metaphase GFP-M2A KI cell. The image is a maximum-intensity projection of two optical planes spanning a 4 µm Z-depth. (E2) An enlargement of the boxed, in-focus region in E1. (E3) The yellow PIV arrows indicate the directions and magnitude that individual M2A filaments within the boxed region in E2 moved in subsequent video frames over 90s (groups of filaments that moved in the apical and basal directions are circled using red and white dashed lines, respectively; see also Movie 10). (F) Cartoon depicting the collective, spatially patterned and bidirectional movements of filaments along the metaphase cell’s apicobasal axis. Scale bars: (B1, C1) 5 µm, (E1) 4 µm, (E2) 2 µm.

In an effort to discern how the myosin cage might achieve dynamic stability while maintaining a rounded shape, we tracked the motions of individual myosin filaments during time lapse movies of metaphase cells. The yellow arrows in Figure 5, Panel E3, show the direction of motion of all the cortical M2A filaments that were in focus within the boxed region of the representative metaphase cell in Panel E1 (this region is enlarged in Panel E2). What is clear is that most if not all of the filaments are moving in groups in either the apical or basal direction (groups moving apically and basally are circled in Panel E3 using red and white dashed lines, respectively; see also Movie 10). We suggest that these collective, spatially pattered and bidirectional movements of filaments along the metaphase cell’s apical/basal axis generate a polarized and differential cortical tension that allows mitotic cells to locally regulate cortical tension so as to maintain a rounded shape in the presence of varying external compressive forces. Key to the success of this mechanism is that both cage organization and collective filament dynamics align with the mitotic cell’s apical/basal axis.

### The myosin filament strings comprising the metaphase cage move laterally during anaphase to help create a contractile ring aligned for planar division

The next major task for M2 is to assemble into a contractile ring. As mentioned in the Introduction, ring creation is thought to involve a combination of de novo actin and myosin filament assembly at the midzone and a cortical flow of preexisting, peripheral, mixed orientation actomyosin assembles that then undergo rearrangement at the midzone [16, 17, 52]. The fact that the M2A filament cage dominates the metaphase cortex suggested to us that it probably contributes to the formation of the contractile ring. Moreover, the fact that the filament strings comprising the cage are parallel to the cell’s apical/basal axis means that they could contribute to the formation of a contractile ring aligned for planar division by simply moving laterally i.e. without the need for large scale rearrangement. Consistent with this thinking, time lapse imaging showed that many of the M2A filament strings that are in focus move into the midzone to help form the contractile ring as the cell progresses through mitosis (Fig. 5, Panels A1-A5 and Movie 11; see the figure legend for details). To support this conclusion, we tracked the motions of individual M2A filaments during contractile ring formation. The yellow arrows in Figure 5, Panel B2, show the direction of motion of all the cortical M2A filaments that were in focus within the boxed region of the representative metaphase cell in Panel A2 (this region is enlarged in Panel B1). What is clear is that the majority of individual filaments present within this region moved in concert toward the midzone of the cell during anaphase to help form the contractile ring. We conclude that the lateral coalescence of the metaphase actomyosin filament strings, which of note is probably guided by the centraspindlin complex so that the coalescing strings are oriented correctly relative to the anaphase spindle, plays a central role in forming a contractile ring designed for planar cell division (see the cartoon in Figure 5, Panel C).

### The fact that the contractile ring created by cage coalescence is anchored at the dividing cells apical surface dictates the basal-to-apical progression of the cleavage furrow

The final major task for M2A is to power contractile ring closure so as to drive cleavage furrow progression. As mentioned in the Introduction, while the contractile ring closes in symmetric fashion in cells dividing in isolation, it closes in asymmetric, basal to apical fashion in cells dividing in within a polarized epithelium [53, 54]. The resulting basal to apical progression of the cleavage furrow serves to allow in coordinated fashion the forming of new cell: cell contacts between the new daughter cells, and to preserve apical tight junctions, essential for barrier function, until the final moments of division [55, 56]. Importantly, the fact that the ends of the cage-derived myosin filament strings present within this contractile ring are anchored at the cell’s apical surface but not at its basal surface should dictate the asymmetric, basal to apical progression of the cleavage furrow. Moreover, this key feature means that the progressing contractile ring should appear as a “half ring” that is continuous at the bottom but not at top because it is anchored at the outer edges of the cell’s apical surface. To confirm this, we imaged the progression of the cleavage furrow, the contractile ring, and the cell’s apical domain in 3D (Fig. 6, Panel D). The YZ views in Figure 6 show the cleavage furrow as it progresses in basal to apical fashion (the arrowheads mark the leading edge of the progressing furrow; the asterisks mark the dividing cell at time 0 and the daughter cells at time 720s; see also Movie 12). As anticipated, the XZ views show that the contractile ring appears as a half ring that progresses in basal to apical fashion until it flattens out against the cell’s apical surface (the white arrowheads in the four images mark the bottom of the half ring and and the red arrows mark its apical attachment sites; the asterisk at time 0 marks the dividing cell). The apical views show that the flattening of the contractile ring against the dividing cell’s apical surface at 720s is accompanied by small lateral indentations of the cell’s apical surface as the new daughter cells appear (compare the positions of the red arrows at 720s to those at 540s; the apical boundary of this dividing cell is outlined with a white dashed line; the asterisk at time 0 marks the dividing cell and the asterisk at 720s marks the two new daughter cells). The graph presents the quantitation of M2A dynamics in this representative dividing cell, with the blue plot showing the decrease in the height of the half ring as the furrow ingresses and the green plot showing the increase in apical M2A signal as the furrow ingresses. Finally, comparing the position over time of the two apical sites where the half ring is anchored in the XZ views with the outline over time of this cell’s apical domain suggests that the half ring is anchored at the cell’s tight junctions (see the cartoon in Fig. 6, Panel F). Consistent with this, 3D lattice light sheet images of dividing cells stained for the tight junction marker ZO-1 show that the two apical ends of the half ring appear anchored to the dividing cell’s tight junctions (Fig. 6, Panels G1-G4).

**Figure 6.**
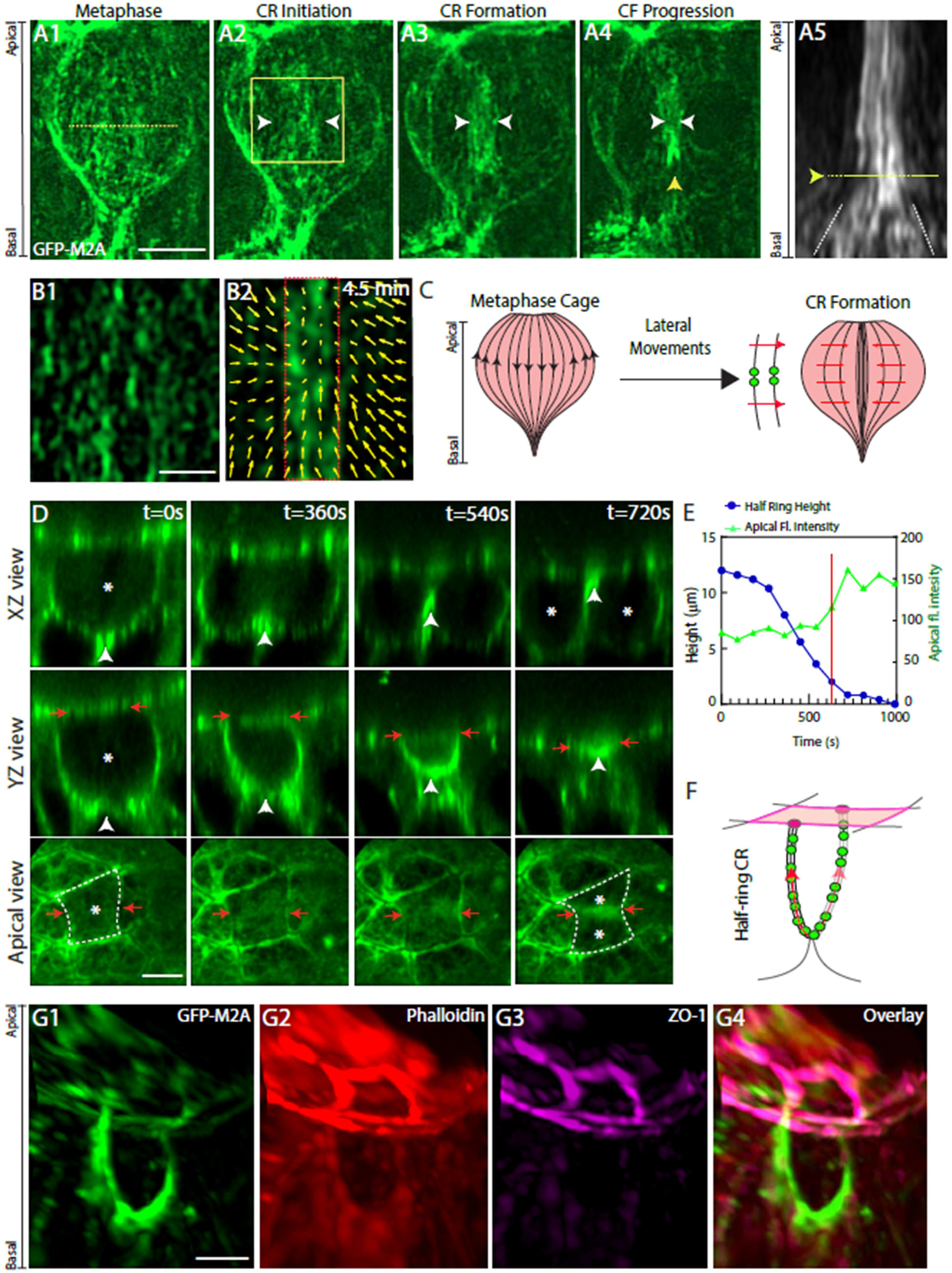
The M2A filament strings comprising the metaphase cage coalesce laterally to help form a contractile ring designed for planar cell division and asymmetric cleavage furrow progression. (A-C) The myosin filament strings comprising the metaphase cage contribute to the formation of a contractile ring aligned for planar cell division. (A1-A4) Snapshots from a representative, AiryScan time lapse movie of a dividing cell within a GFP-M2A KI organoid (single plane) taken at metaphase (A1), the initiation of contractile ring (CR) formation (A2), the formation of the contractile ring (A3), and the initiation of basal-to-apical cleavage furrow (CF) progression (A4) (see also Movie 11). The white arrowheads in A2-A4 bracket the coalescing M2A filament strings, while the yellow arrowhead in A4 marks the initiation of basal-to apical cleavage furrow progression. (A5) Kymograph of M2A filament string coalescence drawn at the dotted line in A1. (B1) Higher magnification image of a portion of a metaphase cage at anaphase onset. (B2) The yellow arrows show the direction of motion of all the individual filaments within the boxed region in B1 over 4.5 minutes as they coalesce to help form the contractile ring (bounded by red dashed lines). (C) Cartoon depicting the contribution that the myosin filament strings comprising the metaphase cage make to forming the contractile ring. (D-G) The anchoring at the dividing cell’s apical surface of the contractile ring created by M2A filament string coalescence dictates basal-to-apical cleavage furrow progression in the form of a progressing “half ring”. (D) XZ views (top row), YZ views (middle row) and apical views (bottom row) of a representative dividing within a GFP-M2A KI organoid at 0, 360, 540, and 720 seconds. The white arrowheads in the XZ and YZ views mark the bottom of the contractile “half ring” as in progresses apically. The red arrows indicate the apical endpoints of the half-ring in the YZ views and the apical boundaries of the dividing cell in the apical views. The single asterisk in the time 0 views and the two asterisks in the 720 second views mark the dividing cell and the two daughter cells, respectively. (E) Complete time course for the changes in half ring height (blue) and apical surface fluorescence intensity (green) for the dividing cell in Panel D. (F) A schematic of the contractile half ring that drives asymmetric cleavage furrow progression in polarized epithelia. (G1-G4) 3D-rendered, Lattice Light Sheet images of a GFP-M2A-labeled contractile half ring (G1) stained with Phalloidin (G2) and for ZO-1 (G3), together with the overlaid image (G4). Scale bars: (A1, D) 5 µm; (B1) 2 µm; (G1) 10 µm.

Taken together, our data argues that the M2A filament strings comprising the metaphase M2A cage undergo three types of concerted motions to support mitosis (Fig. S2). First, they undergo bidirectional, apical/basal motions to support metaphase cell rounding in the presence of variable compressive forces. Second, they undergo lateral and centralizing motions to drive contractile ring formation. Third, they undergo basal to apical motion to drive the asymmetric, basal to apical progression of the cleavage furrow. While the dynamics of individual M2A filaments and filament strings during this last step cannot be resolved optically, we suggest that the M2A filament string dynamics seen during the previous two steps are harnessed to drive this last step. Along those lines, we also suggest that the enormous accumulation of M2A that occurs at the base of the dividing cell to drive INM, and that persists to a significant extent throughout mitosis (see Fig. 3, Movie 4), likely facilitates this third step.

### M2A knockdown leads to a multiple lumen phenotype

To access the significance of the M2A filament cage for driving all aspects of cell division within a polarized epithelium, and to determine the effect of its disruption on epithelial integrity, we transduced GFP-M2A KI organoids expressing the F-actin reporter Td-Tomato LifeAct with a lentivirus harboring a validated M2A-specific shRNA. Successful knockdown was evidenced by an even reduction in GFP fluorescence across the entire organoid. Figure 4, Panel A1, shows the GFP-M2A signal in one such organoid immediately above a non-transduced, control organoid. Comparing the LifeAct signals in these two organoids (Fig. 7, Panel A2) reveals a dramatic difference in morphology, with the control organoid exhibiting a single lumen and the knockdown organoid appearing multilobed. Comparing knockdown organoids exhibiting differences in GFP signal strength showed that the multi-lobed phenotype scales with the degree of knockdown: the dimmer the GFP-M2A signal, the more lobes the organoid exhibits (Fig. 6, Panels B1, B2, C1, C2, D1, D2). Importantly, staining for the apical domain marker ZO-1 showed that each of these lobes corresponds to a separate lumen (Fig. 7, Panel E). Of note, measurements of fluorescence intensity per unit area indicated that maximal knockdown was ∼70%, presumably because some amount of M2A is required for viability. Together, these results show that M2A plays an important role in maintaining lumen integrity.

**Figure 7:**
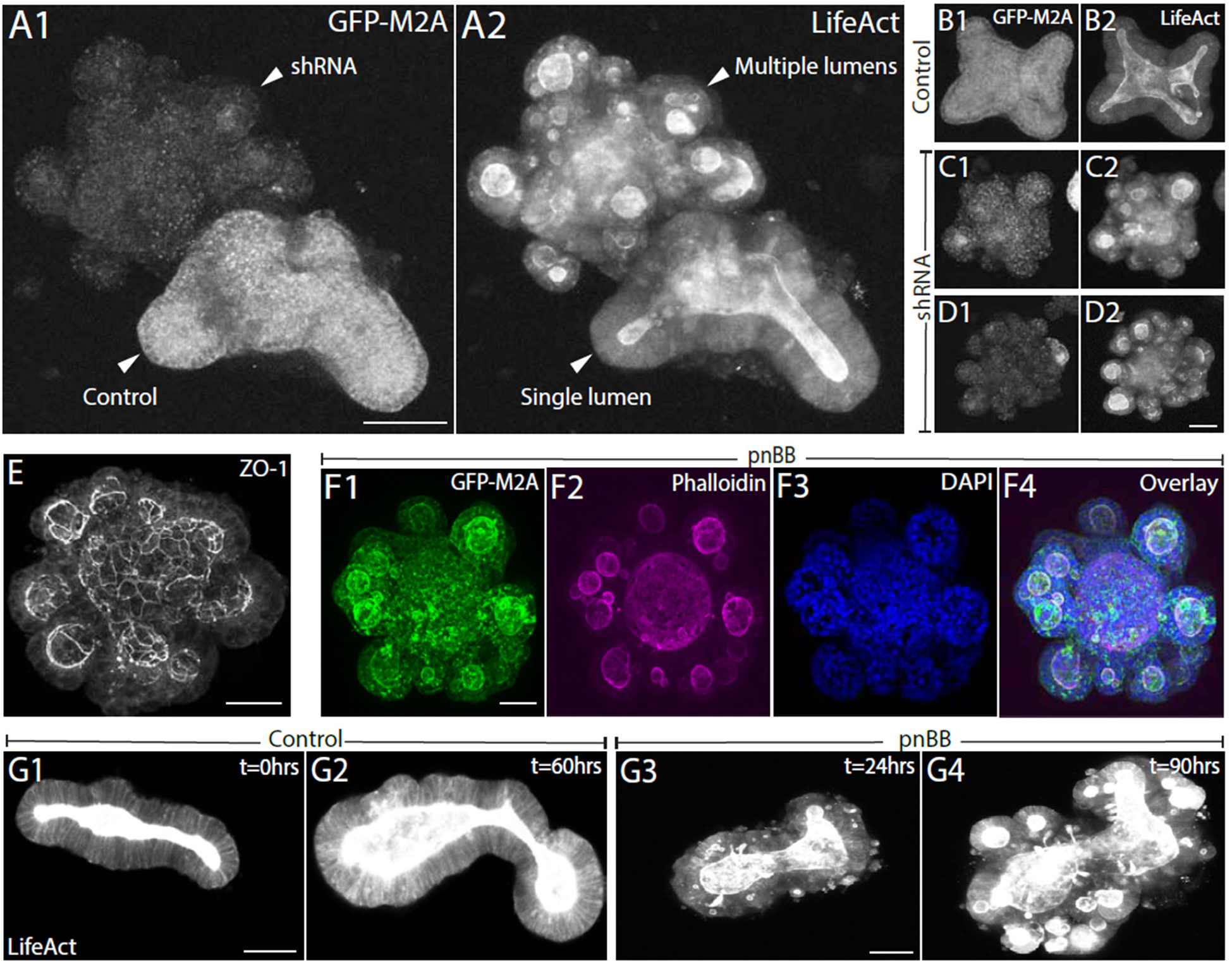
Attenuation of M2A function using RNAi or Blebbistatin leads to a multiple lumen phenotype. (A1) Shown are representative examples of a control GFP-M2A KI organoid next to a GFP-M2A KI organoid that had been transduced with a lenti virus driving the expression of an shRNA against M2A (note the loss of GFP signal; fluorescence intensity measurements are consistent with an ∼70% knockdown of M2A in such organoids). (A2) The signal for F-actin in these same two organoids, which were also expressing Td-Tomato LifeAct to highlight the transition from a single lumen to what looks like multiple lumens when M2A function is attenuated. (B-D) GFP-M2A signals (B1-D1) and Td-Tomato LifeAct signals (B2-D2) for a control GFP-M2A KI organoid (B1, B2), a M2A knockdown organoid where the knockdown was moderate (C1, C2), and a M2A knockdown organoid where the knockdown was more penetrant (D1, D2). (E) Representative, maximum intensity, projected AiryScan image of a M2A knockdown organoid stained for ZO-1. (F1-F4) Representative, maximum intensity, projected AiryScan images of a GFP-M2A KI organoid that was treated with 30 µm pnBB for 72 hours (F1) followed by staining with Phalloidin (F2) and DAPi (F3). (F4) The overlay of the images in F1-F3. (G1-G4) Stills from representative time lapse movies of a control Td-Tomato LifeAct expressing organoid at 0 (G1) and 60 hours (G2) and a pnBB-treated, Td-Tomato LifeAct expressing organoid at 24 (G3) and 60 hours (G4). See also Movie 13. Scale bars: (A1, D2, E) 20 μm; (F1, G1, G3) 30 μm.

### Paranitro-blebbistatin treatment phenocopies M2A knockdown

Growing organoids in the presence of the M2 inhibitor paranitro-blebbistatin (pnBB; 30 µM) led to the formation of multiple lumen structures that were indistinguishable from M2A knockdown organoids (Fig. 7, Panels F1-F4). Most revealing is time lapse imaging, where control organoids can be seen to maintain a single lumen as they grow (Fig. 7, Panels G1 and G2) while pnBB-treated organoids generate multiple lumens (Fig. 7, Panels G3 and G4; see also Movie 13). Of note, the maximal inhibition of M2A’s actin-activated ATPase activity obtainable *in vitro* using pnBB is ∼60% (reached at a concentration of ∼40 µM) [57]. Like shRNA-mediated knockdown of M2A, therefore, a significant amount of M2A activity remains in pnBB-treated organoids, and this residual activity is likely required for viability. Consistently, organoids cultured from the M2A conditional knockout mouse died within three days following treatment with a lenti virus expressing cre recombinase (Fig. S3). Interestingly, restoration of M2A contractility by pnBB washout, if done before the multiple lumen phenotype progresses too far, results in lobe merger to restore the single lumen (Movie 14).

### M2C does not contribute to the maintenance of lumen integrity

While pnBB inhibits M2C as well as M2A [57], its effect on lumen integrity is specific for M2A, as organoids made from the M2C knockout mouse maintain a single lumen as they grow (Fig. S4, Panel A). Consistently, inhibition of M2A in M2C KO organoids using pnBB yields the multi-lumen phenotype (Fig. S4, Panel B). This result also argues that the multi-lumen phenotype exhibited by M2A-shRNA knockdown organoids is not due to an imbalance in apical versus basal tension caused by the persistence of M2C-dependent apical tension when M2A-dependent basal tension is attenuated.

### pnBB-treated organoids exhibit orthogonal cell divisions

To investigate the possibility that the multi-lumen phenotype arises from a defect in cell division, we imaged the orientation and motion of DNA in control and pnBB-treated cells expressing RFP-Histone 2B. As expected, control organoids exhibit planar cell divisions in which the duplicated chromosomes move parallel to the plane of the epithelium during anaphase (Fig. 8, Panel A1 and Movie 15). Consistently, tubulin staining shows that the mitotic spindle in these cells is parallel to the epithelial plane (Fig. 8, Panel A2), and expressing RFP-neuromodulin to mark plasma membranes shows that the daughter cells end up side-by-side in the epithelial plane (Fig. 8, Panels A3 and A4). In contrast, pnBB-treated organoids exhibit orthogonal divisions in which the duplicated chromosomes move perpendicular to the plane of the epithelium during anaphase (Fig. 8, Panel B1 and Movie 16). Consistently, the mitotic spindle in these cells is now perpendicular to the epithelial plane (Fig. 8, Panel B2), and the daughter cells end up being stacked one on top of the other rather than being side-by-side (Fig. 8, Panels B3 and B4). Movie 18 shows several such events (marked by arrowheads) during a 10 hour time lapse movie of a pnBB-treated organoid expressing RFP-neuromodulin.

**Figure 8:**
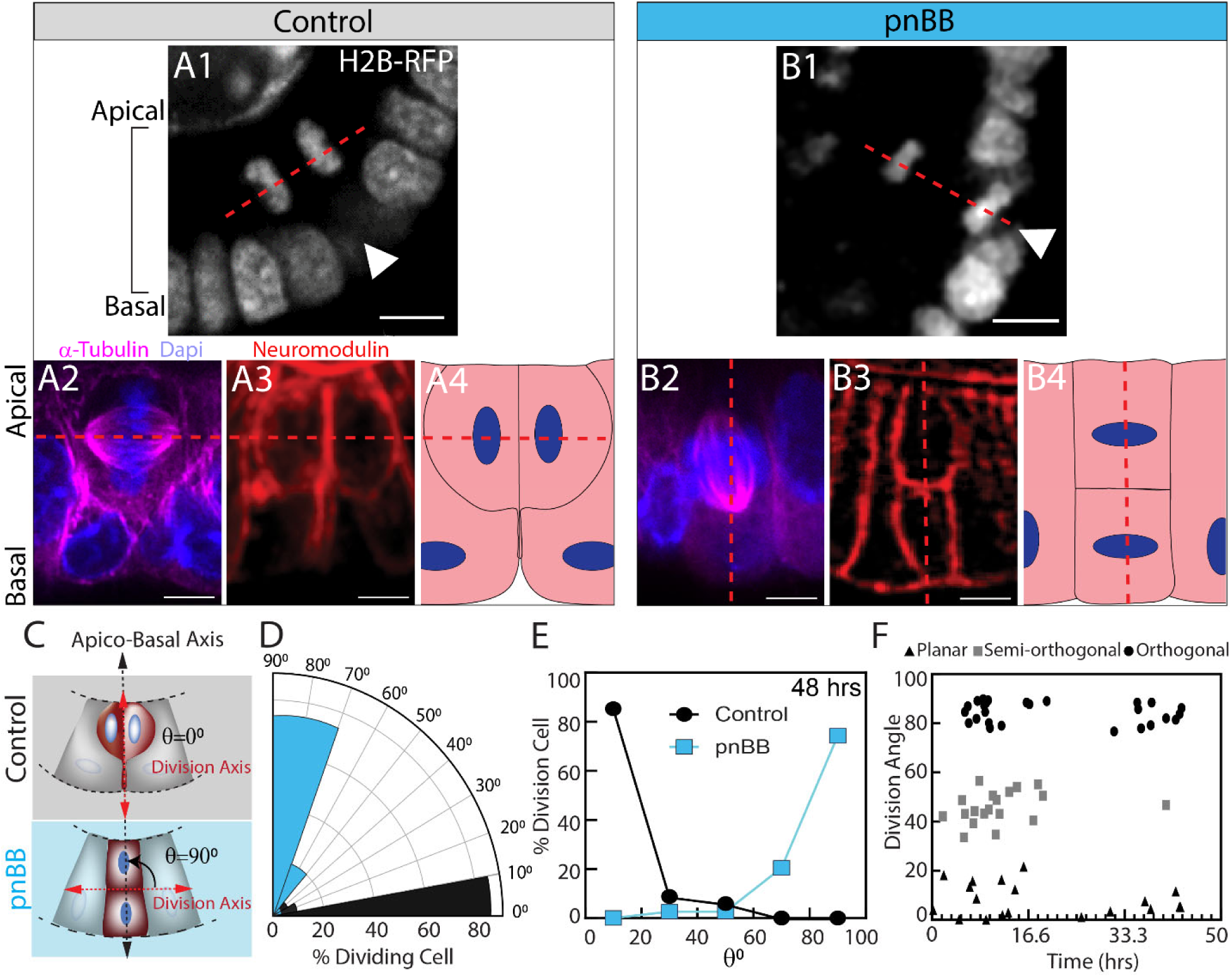
Attenuation of M2A function shifts the division plane from planar to orthogonal. (A1) Single frame from a time lapse movie of a dividing cell at late anaphase (arrowhead) within a control, H2B-RFP-expressing organoid. See also Movie 15. (A2) A metaphase cell within a control, RFP-neuromodulin-expressing organoid that was fixed and stained with anti-α-tubulin and DAPI. (A3) The neuromodulin signal immediately after completion of cytokinesis. (A4) Cartoon depicting the arrangement of daughter cells immediately after completion of a normal, planar cell division. The red dashed lines in A1-A4 mark the division axis. (B1-B4) Same as in A1-A4 except that the dividing cell within this organoid was treated with pnBB (B4 shows a cartoon depicting the arrangement of daughter cells immediately after completion of an orthogonal cell division) (See Movie 16). (C) Cartoon depicting the cell division axis (red line) relative to the cell’s apico-basal axis (black line) in a control organoid, where theta (θ) approaches O°, and a pnBB-treated organoid, where θ approaches 90°. (D) Distribution of division angles for control (black) and pnBB-treated (blue) organoids at 48 hours (from 30 dividing cells in 3 organoids for each condition). (E) Distribution of division angles for control (black) and pnBB-treated (blue) organoids at 48 hours. (F) The distribution of division angles over time following addition of 30 µM pnBB, binned as planar (0° to 30°, triangles), semi-orthogonal (30° to 60°, squares), and orthogonal (60° to 90°, circles). Scale bars: 10 µm.

Quantitation of division angles in time lapse movies of neuromodulin-marked organoids, where an angle of 0° corresponds to a perfectly planar division and an angle of 90° corresponds to a perfectly orthogonal division (Figure 8, Panel C), showed that ∼85% of cells in control organoids divide within 10° of perfectly planar while ∼80% of cells in pnBB-treated organoids divided within 25° of perfectly orthogonal (Fig. 8, Panels D and E). Scoring the division angle at increasing times after pnBB addition showed that the proportion of cells exhibiting planar divisions (0° to 30°), semi-orthogonal divisions (30° to 70°), and orthogonal divisions (70° to 90°) shifts progressively to orthogonal divisions over time (Fig. 8, Panel F). Together, these data show that attenuating M2A function results in a dramatic shift in the cell division axis from planar to orthogonal.

### pnBB treatment inhibits both INM and metaphase cell rounding

To determine what causes the division plane to shift from planar to orthogonal when M2A function is attenuated, we measured the magnitudes of INM and metaphase cell rounding in control and pnBB-treated organoids given that both events are known to be myosin 2 dependent ([37, 58]; see also Figure 3). With regard to INM, Figure 9, Panels A1 to A8, show single frames from representative time-lapse movies of dividing cells in control (A1-A4) and pnBB-treated (A5-A8) organoids expressing RFP-Histone 2B to mark nuclei/chromatin. Measurements of the distance traveled by the nucleus/chromatin from its pre-mitosis position at the cell base (marked with an asterisk) to its more apical position at metaphase (marked with a dot) showed that the nucleus/chromatin in pnBB-treated cells travels in the apical direction about half as far as in control cells (Fig. 9, Panel B; note that the distance traveled was normalized to the height of the corresponding surrounding epithelium). Consistent with this reduction in apical travel, the height of the upper daughter cell in the daughter cell stack formed by an orthogonal division is on average significantly larger than the height of the lower daughter cell (Fig. 9, Panel C). The M2A-dependent pushing force driving INM also creates a pulling force that invaginates the cell’s basal surface at a site where FAK-positive focal adhesions are concentrated (Fig. 9, Panels D1 and E1-E4, Movie 17). Consistent with this, and with the observed defect in INM, metaphase cells in pnBB-treated organoids generate much smaller basal invaginations than metaphase cells in control organoids (Fig. 9, Panels D2 and D3).

**Figure 9:**
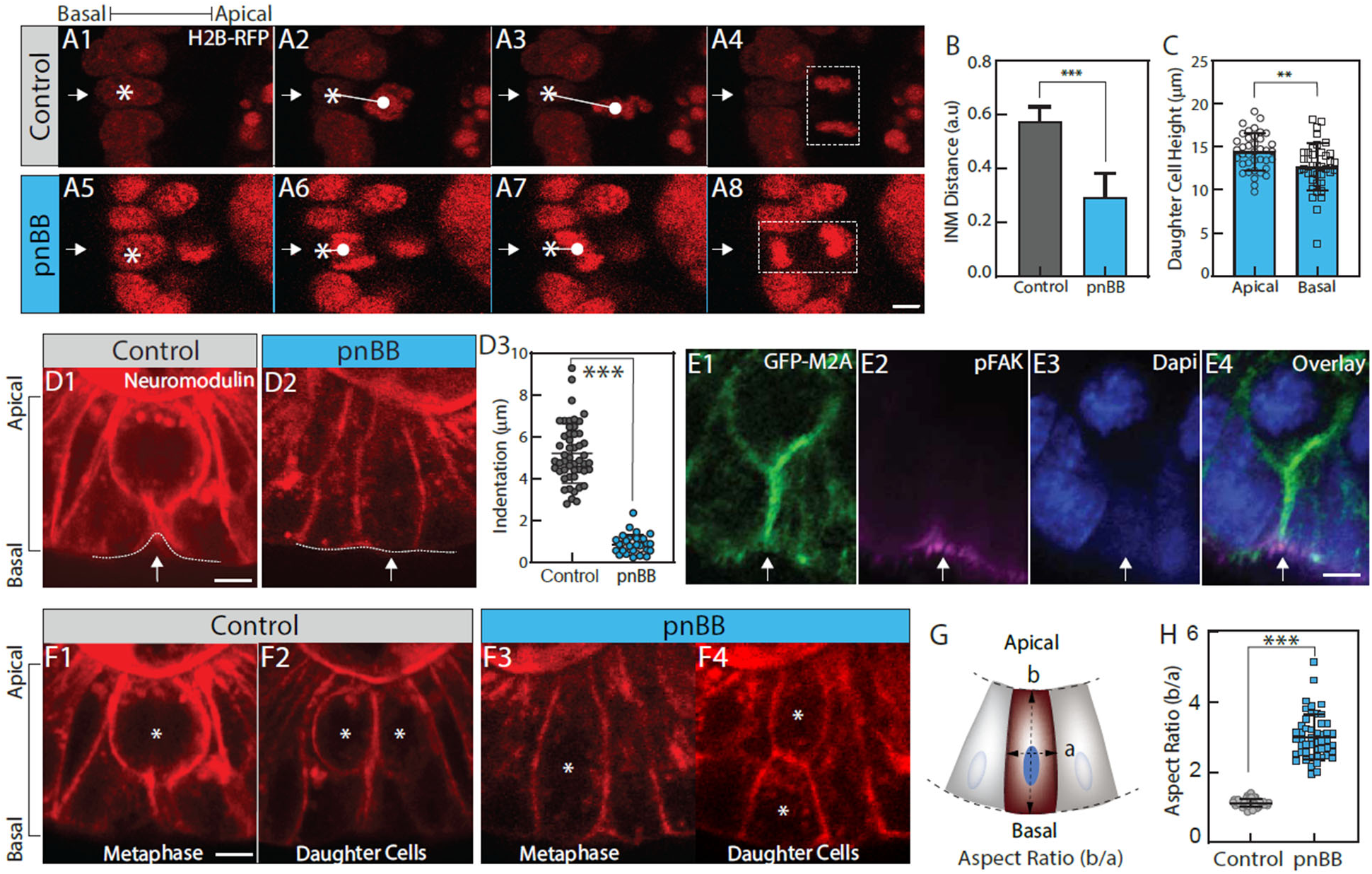
Attenuation of M2A function impairs INM, basal surface indentation, and mitotic cell rounding. (A1-A4) Stills from a representative, time lapse, AiryScan movie of a dividing cell undergoing INM within a control, H2B-RFP-expressing organoid. The arrow in A1-A4 marks the dividing cell, and the asterisk and dot in A1-A3 mark the initial position of the nucleus and the position of the chromatin at metaphase, respectively. The line connecting the asterisk and dot represents the distance the nucleus/DNA traveled during INM. (A4) The distribution of chromatin in this control cell at late anaphase, indicating a planar division. (B1-B4) As in A1-A4 except that this organoid was treated with pnBB (B4 shows the distribution of chromatin in this cell at late anaphase, indicating an orthogonal division). (B) Quantification of the distance the nucleus traveled during INM in control and pnBB-treated organoids, normalized for the width of the epithelium (n = 40 cells for both conditions). (C) Quantification of the heights of the upper and lower daughter cells created by orthogonal divisions (n = 40 for both conditions). (D1) Representative image of a control cell at metaphase within an RFP-Neuromodulin expressing organoid (the dashed white line and arrow indicate the indentation of the cell’s basal surface; see also Movie 17). (D2) As in D1 except that this metaphase cell was within a pnBB treated organoid (see also Movie 18). (D3) The length of basal indentations for control and pnBB treated cells at metaphase (n = 50 for both conditions). (E1-E3) Representative Airyscan images of a control, metaphase cell (marked with an asterisk) within a GFP-M2A KI organoid (E1) that was fixed and stained with anti-pFAK (E2) and DAPI (E3). (E4) Overlay of E1-E3. (F1) Representative AiryScan image of a control cell at metaphase within an RFP-Neuromodulin-expressing organoid. (F2) This same cell just after completion of cytokinesis (the two daughter cells are marked with asterisks). (F3 and F4) As in F1 and F2 except that this dividing cell was within a pnBB treated organoid. (G) Cartoon depicting how the dividing cell’s short (a) and long (b) axes were measured at metaphase to calculate the cell’s aspect ratio (b/a). (H) Aspect ratios of metaphase cells in control and pnBB-treated organoids (n = 50 cells for both conditions). Scale bars: 5 µm.

With regard to metaphase cell rounding, Panels F1 and F2 in Figure 9 show single frames from a representative time-lapse movie of a metaphase cell (F1) and its subsequent daughter cells (F2) in a control organoid, while Panels F3 and F4 show the same for a pnBB-treated organoid (the label is neuromodulin-RFP, and the metaphase cell and it daughters are marked with asterisks; see also Movie 18). Measurements of metaphase cell aspect ratio (longer axis(b)/shorter axis (a); see Panel G) showed that control cells are on average close to being perfectly round (aspect ratio 1.1 ± 0.2) while pnBB-treated cells are on average significantly elongated along the apical/basal axis (aspect ratio 2.8 ± 0.7) (Fig. 9, Panel H). Together, these data argue that, as in fly epithelia [57], the geometric defects imposed by impaired INM and metaphase cell rounding drive a 90 degree rotation of the spindle so that M2A-inhibited cells can divide along their long axis. Finally, to link the metaphase M2A filament cage, which we argue drives metaphase rounding, to the effect of M2A inhibition on metaphase rounding, we imaged the cage in rounded mitotic cells within 2 h of treatment with 30 µM pnBB (i.e. before mitotic rounding was completely lost). Movie 19 shows that in such cells the M2A filament cage undergoes progressive disassembly that is accompanied by a transition in cell shape from rounded to elongated.

### Orthogonal divisions generate nascent lumens that then mature into independent lumens

In an effort to link orthogonal divisions to the initiation of supernumerary lumen formation, we stained GFP-M2A KI organoids with the apical domain marker ZO-1 eight hours after adding pnBB. This approach revealed small circular signals for ZO-1 where newly formed stacked daughter cells abut (Fig. 10, Panels A2 and A5; the dotted circles in A5 mark the nuclei of the stacked daughter cells created by an orthogonal division). These ZO-1 positive circles are also enriched for GFP-M2A and F-actin (Fig. 10, Panels A1 and A3), and they are prominent in similarly-treated organoids stained for F-actin and ZO-1 and imaged in 3D by Lattice Light Sheet microscopy (Fig.10, Panels B1 and B2; see also Movie 20).

**Figure 10:**
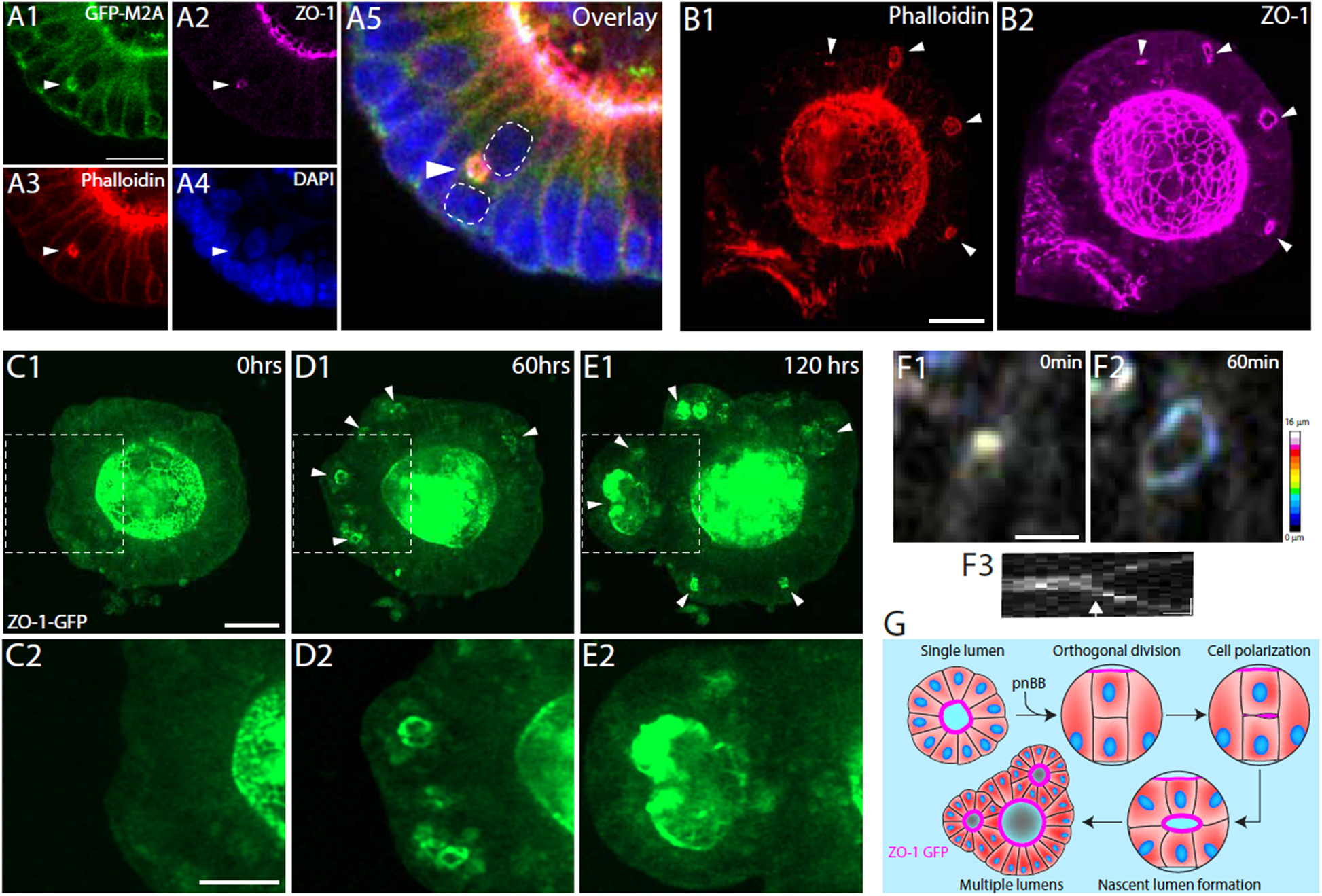
Nascent lumens that originate at the interface between the stacked daughter cells generated by an orthogonal division grow into mature lumens. (A1-A4) Airyscan images of a portion of a representative GFP-M2A KI organoid that had been treated with 30 μM pnBB for 24 hours (A1) and then fixed and stained with anti-ZO-1 (A2), Phalloidin (A3), and DAPi (A4). (A5) Overlay of A1-A4 (the dotted lines encircle the two stacked daughter cells arising from an orthogonal division). (B1, B2) 3D rendered, Lattice Light Sheet images of an organoid treated with 30 μM pnBB for 8 hours and then fixed and stained with Phalloidin (B1) and anti-ZO-1 (B2) (see also Movie 20). (C-E) Stills from a representative, time lapse AiryScan movie of a GFP-ZO-1 KI organoid taken at 0 hours (time of paBB addition) (C1), 60 hours (D1), and 120 hours (E1). The boxed areas are enlarged in (C2), (D2), and (E2). See also Movie 21. (F) Shown is a depth coded example of polarity establishment following an orthogonal division with the initial polarization of the orthogonal daughter cell (F1), the formation and expansion of a nascent lumen (F2), and a kymograph derived from the time-lapse movie (F3). (G) Schematic summarizing the pathway by which an orthogonal division leads to polarity establishment in the stacked daughter cells, then nascent lumen formation, and finally the emergence of the multiple lumens. Scale bars: (A, B, C, D) 20 µm; (F1) 5 µm.

To confirm that these ZO-1-positive circles correspond to nascent lumens that subsequently mature into the numerous, independent lumens seen in ZO-1 stained organoids several days after pnBB addition, we performed time lapse imaging of organoids made from a GFP-ZO-1 KI mouse [59] to follow apical domains from lumen inception to supernumerary lumen formation. Consistent with the still images in Figure 10, Panel E, dynamic imaging of pnBB-treated GFP-ZO-1 KI organoids shows that the nascent lumens formed at sites of orthogonal division do indeed transform into independent, mature lumens (Fig. 10, Panels C1, C2, D1, D2, E1, E2; see also Movie 21; Panels F1 to F3 show the initial expansion of a single nascent lumen; Panel G, cartoon). Close inspection of Movie 21 shows that while most lumens grow in isolation, some also enlarge by fusing together.

## DISCUSSION

Cell division within epithelia presents a unique mechanical challenge because the cells must undergo extensive shape changes to achieve mitotic rounding while remaining integrated within a mechanically constrained tissue [44]. This is particularly relevant for the intestinal epithelium, where billions of cell divisions occur each day to maintain tissue homeostasis [60-62]. Although actomyosin contractility is known to contribute to mitotic rounding and cell division, how it supports these processes within a physiologically relevant polarized epithelial environment has remained largely unexplored. Here, using intestinal organoids derived from a GFP-M2A knockin mouse, together with super-resolution microscopy, pharmacological perturbations, and laser ablation, we identify a previously unrecognized metaphase M2A filament structure that coordinates multiple aspects of epithelial mitosis, including metaphase rounding, division orientation, and cleavage furrow progression, thereby ensuring faithful epithelial cell division and tissue homeostasis

While interphase cells display M2A enrichment along basal stress fibers and lateral membranes, metaphase cells in both organoids and tissue exhibit a strikingly different organization in which M2A assembles into a cortical, cage-like network of bipolar filaments that are aligned along the dividing cell’s apical-basal axis. Thus, entry into mitosis within epithelia is accompanied by a transition from a relatively disordered cortical organization to a highly ordered cortical apico-basal architecture. Notably, this organization contrasts sharply with the isotropic and randomly oriented cortical distribution of myosin filaments observed in isolated mitotic cells, both in our experiments and in a large number of previous studies (reviewed in [15, 18, 63-66]). These findings argue that polarized epithelial cells create a unique contractile structure that allows them to perform those aspects of cell division unique to their situation: metaphase rounding in the context of a mechanically constrained tissue environment, a planar axis of division, and asymmetric, basal to apical cleavage furrow progression.

We present evidence that the M2A filament cage performs an important mechanical function during metaphase rounding that extends beyond the established role of cortical myosin as a source of tension opposing intracellular hydrostatic pressure. Laser ablation on one side of the metaphase cortex consistently induced displacement away from the ablation site together with local shape deformation, indicating that the cage contributes to force balance across the mitotic cortex. If cortical tension functioned primarily to counteract hydrostatic pressure, local loss of tension would be expected to promote expansion toward the ablated side [20, 67]. Instead, the observed displacement away from the ablation site and the reduction in local curvature indicates that the ablated region is dominated by external compressive forces. These observations argue that the metaphase M2A filament cage is not merely a contractile shell opposing hydrostatic pressure, but rather a specialized contractile structure that distributes and balances forces across the metaphase cortex to resist tissue-derived compressive forces. These findings prompted us to examine M2A filament cage dynamics in greater detail. PIV analysis of super-resolution time-lapse images revealed spatially patterned, bidirectional cortical flows of filaments that were largely parallel to the cell’s apical-basal axis and that occurred throughout metaphase. These observations argue that robust mitotic rounding in epithelia is driven by dynamic apico-basal cortical contractility rather than by uniformly distributed cortical contractility. Together, these findings support a model in which the M2A filament cage functions as a specialized force-balancing structure that spatially organizes cortical contractility to promote isotropic mitotic rounding while resisting tissue-derived compressive forces. In this way, the cage couples mitotic rounding to the mechanical constraints of the surrounding epithelium.

Another defining feature of the metaphase M2A filament cage is that it is not a transient structure that disappears at anaphase onset. Instead, this preassembled contractile scaffold repositions toward the spindle midzone and progressively coalesces to help form the contractile ring. This mechanism of contractile ring formation may provide distinct mechanical advantages in three-dimensional epithelial tissues, where polarity, geometric confinement, and mechanical coupling to neighboring cells impose additional constraints on cytokinesis. Classical models of contractile ring formation propose that spindle-dependent RhoA signaling specifies an equatorial zone where actomyosin is assembled de novo and then reinforced by flows of isotopically-distributed cortical actomyosin that are realigned at the midzone through compression and myosin contractility [16, 66]. While our findings do not exclude some contributions from these pathways, they argue that epithelial cells superimpose on them the harnessing of a preassembled metaphase actomyosin structure for contractile ring formation. Importantly, the fact that the M2A filament strings comprising this metaphase structure are parallel to the cell’s apical/basal axis means that they will contribute to the formation of a contractile ring aligned for planar division by simply coalescing laterally, i.e. without the need for either large scale de novo assembly or realignment at the midzone.

A defining feature of cytokinesis in epithelia is that the cleavage furrow progresses asymmetrically from base to apex rather than closing symmetrically as in cells dividing in isolation. This feature serves to maintain barrier function throughout division by restricting the final resolution of the daughter cells until the cleavage furrow has reached the dividing cell’s tight junctions, the main drivers of barrier function. Asymmetric cleavage furrow progression also allows in coordinated fashion the formation of new junctions between the forming daughter cells [43]. Importantly, the fact that the contractile ring in epithelial cells is derived from the metaphase M2A filament cage, together with the fact that the ends of the cage-derived myosin filament strings comprising this contractile ring are not anchored basally but are anchored apically at the cell’s tight junctions, provides a simple mechanical explanation for asymmetric, basal-to-apical cleavage furrow progression. This mechanism also explains why the contractile ring appears as a half ring that progresses base to apex rather than as a complete ring that closes symmetrically. Thus, the metaphase M2A filament cage serves two sequential mechanical functions: it first spatially organizes cortical contractility to maintain mitotic rounding in the presence of tissue-derived compressive forces, and it subsequently remodels into a contractile half-ring that dictates both a planar axis of division and asymmetric cleavage furrow progression.

The physiological importance of the M2A filament cage became evident following either partial depletion of M2A or attenuation of its contractility, both of which resulted in organoids possessing multiple lumens. Mechanistically, disruption of the cage impaired both INM and mitotic rounding, resulting in a shift in division orientation from planar to orthogonal. This shift results in the daughter cells becoming stacked along the epithelium’s apico-basal axis, creating a geometry that promotes *de novo* lumen formation at their interface. Martin and colleagues [58] showed previously that inhibiting myosin 2 function in *Drosophila* epithelial monolayers impairs both INM and metaphase cell rounding, and that these geometrical defects shifted the division plane from planar to orthogonal despite the persistence of functioning polarity cues. While the immediate consequences of M2A inhibition in organoids mirror those findings, they differ from the consequences of orthogonal divisions within *Drosophila* tissues, where the misoriented daughter cells are either extruded or reintegrated into the epithelial monolayer [68-70]. In contrast, dynamic imaging of GFP-ZO-1 KI organoids treated with BB showed that the stacked daughter cells arising from orthogonal divisions remain in place, establish apico-basal polarity, and initiate the formation of independent lumens. These findings demonstrate that accurate division orientation and lumen maintenance emerge not only from polarity cues but also from the mechanical processes that govern mitosis within epithelial tissues.

Previous studies have emphasized the roles played by polarity cues and junctional signaling in maintaining epithelial topology [71-79]. Our findings suggest that these regulatory systems operate in concert with a specialized, M2A-based mitotic structure. By coordinating mitotic rounding, division orientation, and cytokinesis, this mitotic structure links cell-scale force generation to tissue-scale morphogenesis. More broadly, our findings establish that epithelial cell division is governed not simply by the magnitude of actomyosin-based contractility, but by its spatiotemporal organization into a specialized cortical structure that functions to preserve epithelial topology.

## METHODS

### Mouse strains

All animal experiments were approved by the Animal Care and Use Committee of the National Heart, Lung, and Blood Institute, in accordance with the National Institutes of Health guidelines. The animal colony was maintained under pathogen-free conditions. All procedures were conducted under an animal research protocol in accordance with the National Institutes of Health Animal Care and Use Guidelines. Adult mice between 8 and 16 weeks of age were used for all experiments. Both male and female mice were included unless otherwise specified. Animals were euthanized prior to tissue collection according to approved protocols. The following mice were used in this study: C57BL/6 (WT), C57BL/6 GFP-Myosin 2A knockin (GFP-M2A KI), C57BL/6 GFP-Myosin 2B knockin (GFP-M2B KI), C57BL/6 GFP-Myosin 2C knockin (GFP-M2C KI), C57BL/6 Myosin 2C knockout (M2C KO), C57BL/6 Myosin 2A conditional knockout (M2A cKO), and C57BL6 GFP-ZO-1 knockin (GFP-ZO-1 KI). The GFP-M2 KI mice and the M2A cKO mouse were gifts of the late Robert S Adelstein (NHLBI, NIH) except that we backcrossed them to C57BL/6, the M2C-KO was created on the C57BL/6 background using CRISPR (unpublished), and the GFP-ZO-1 KI mouse was a gift of Terry Lechler (Duke University) except that we backcrossed it to C57BL/6.

### Isolating Crypts from KI and KO mouse intestines

Mouse small intestines were isolated from 8 to 10 week old GFP M2 KI mice, the M2C KO mouse, or the M2A cKO mouse following euthanasia using an NIH-approved protocol. Intestinal contents were removed by thoroughly flushing the lumen with cold PBS, and visible connective tissue was removed. Intestines were opened longitudinally, washed extensively with cold PBS, cut into 2 to 4 mm fragments, and the issue fragments washed repeatedly with cold PBS until the supernatant was clear. For crypt isolation, tissue fragments were incubated in Gentle Cell Dissociation Reagent (STEMCELL Technologies, #100-0485) for 20 min at room temperature with gentle rocking. Crypts were subsequently released by serial trituration in 0.1% BSA in PBS and collected as four sequential 10 mL fractions. Each fraction was filtered through a 70 µm cell strainer and centrifuged at 300 × g for 5 min. Pellets from the third and fourth fractions were resuspended separately in DMEM/F12 media (Gibco, #11320033) and centrifuged again at 300 × g for 5 min. The crypt pellet from either the third or fourth fraction (chosen by visible observation) was resuspended in IntestiCult medium (STEMCELL Technologies, #06005) and growth factor-reduced Matrigel (Corning, #356231) and plated as 50 µL domes in 24 well plates. Matrigel domes were allowed to polymerize at 37°C for 30 min, after which 600 µL of IntestiCult medium containing antibiotics was added to each well.

### Organoid growth and passaging

Organoids were passaged every 5 to 7 days. Briefly, the culture medium was removed, the Matrigel-embedded organoids were suspended in Gentle Cell Dissociation Reagent, and repeated pipetting was used to disrupt the Matrigel and fragment the organoids. The fragmented organoid suspensions were transferred to tubes pre-wetted with 0.1% BSA in PBS and gently rocked at room temperature for 10 to 15 min. Following centrifugation at 300 × g for 5 min at 4°C, the pellets were resuspended in DMEM/F12 and centrifuged again. The resulting pellets were resuspended in IntestiCult medium at the desired split ratio, mixed 1:1 with ice-cold Matrigel, and plated as 50 µL domes in prewarmed 24 well plates or imaging chambers. After Matrigel polymerization at 37°C, cultures were overlaid with IntestiCult medium and maintained with media changes every 2 to 3 days.

### Lentivirus packaging and organoid transduction

Second-generation lentiviral particles were generated using the packaging plasmid psPAX2 (Addgene, #12260) for pLV-RFP-H2B (Addgene, #26001), pLenti-Lifeact-tdTomato (Plasmid #64048), and pLVX-EF1α-Neuromodulin-IRES-Puromycin (Addgene, #134666). Lentivirus production and concentration were performed as previously described (Yim et al., MBoC, 2024)[80]. For stable transduction, mature organoids were partially dissociated into small multicellular crypt fragments using Gentle Cell Dissociation Reagent. Organoids were disrupted by trituration, incubated in Gentle Cell Dissociation Reagent for 15 min at room temperature, and triturated again. Organoid fragments were pelleted at 300 × g for 5 min, washed with cold DMEM/F12, centrifuged again at 300 × g for 5 min, and resuspended in 500 µL IntestiCult medium supplemented with 10 µM Y-27632 ROCK inhibitor (Cayman Chemical, #10005583). Prior to transduction, 4 well imaging chambers were coated with 250 µL per well of Matrigel diluted 1: 50 in PBS and incubated at 37°C for at least 2 hr. The diluted Matrigel was removed immediately before plating the organoid fragments. For each well, the 500 µL organoid suspension was combined with 10 µL of concentrated lentivirus and 0.5 µL of polybrene (Sigma-Aldrich, #TR-1003), with Y-27632 maintained at 10 µM, and the transduction mixture was added dropwise into each coated wells. Cultures were incubated with lentivirus for 6 to 24 hr. The following day, the lentiviral medium was carefully removed, and the cultures were gently washed to remove non-adherent and dead cells. Adherent organoid colonies were overlaid with 300 µL of a 1:1 mixture of growth factor-reduced Matrigel and IntestiCult medium per well. After Matrigel polymerization, 400 µL of IntestiCult medium was added, and the organoids were maintained with media changes every 2 days. Transduced organoids were passaged 4 to 6 days after transduction and, when appropriate, subjected to antibiotic selection to enrich for stably transduced cells.

### Production of MSCV-Cre-GFP retrovirus

For retrovirus production, a transfer plasmid encoding Cre recombinase (Addgene, #24064) was co-transfected with the pCL-Eco helper plasmid (Addgene, #12371) at a 1:1 ratio into HEK293T/17 cells at ∼80% confluency. Viral supernatants were harvested at 24 and 48 hrs post-transfection, pooled, and filtered through a 0.45 µm PVDF filter. Organoids derived from floxed myosin-2 and WT mice were cultured for 24 hrs and then treated with MSCV-Cre-GFP retrovirus to induce myosin-2 deletion. Organoids were imaged 48 to 72 hrs after treatment, with GFP fluorescence used to identify Cre-expressing cells.

### Immunoblotting

To generate lysates for Western blotting, organoids were collected in cold PBS and pelleted by centrifugation at 300 × g for 5 min. Pellets were resuspended 1X Cell Lysis Buffer (Cell Signaling, #9803) that contained a protease/phosphatase inhibitor cocktail (Cell Signaling, #5872). Samples were resolved on a 4 to 12% gradient or 6% NuPAGE Bis-Tris gel (Thermo Fisher Scientific, #XP04120BOX or #XP00060BOX) and transferred onto nitrocellulose membranes (Bio-Rad, #162-0251) using Bio-Rad Trans Blot SD Semi Dry Transfer Cell a semidry transfer system (Bio-Rad, #1703940). Nitrocellulose membranes were blocked in TBST (10 mM Tris (pH 8.0), 150 mM NaCl, and 0.02% Tween 20) supplemented with 5% milk for 2 h, incubated with anti-M2A (Cell Signaling, #3403), anti-M2B (BioLegend, #PRB-445P), or anti-M2C (ProteinTech, #66825-1-Ig) primary antibodies overnight at 4°C, washed 3X with TBST, and incubated with secondary antibody (Jackson ImmunoResearch, #115-035-045) at RT for 2 h. Antibodies were diluted in TBST containing 5% milk. Anti-HSC70 (Santa Cruz Biotechnology, #sc-137239) was used as the loading control. Proteins were detected using SuperSignal West Pico Plus Chemiluminescent Substrate (Thermo Fisher Scientific, #34075) and quantitated using an Amersham Imager 600 (GE Healthcare Life Sciences).

### Organoid fixation and staining

Organoids were collected from Matrigel domes using cold PBS and fixed in 4% paraformaldehyde (PFA) for 30 min with gentle rocking. Following fixation, organoids were washed with PBS and then permeabilized and blocked for 2 h in 5% serum containing 1% Triton X-100. Samples were subsequently washed with immunofluorescence (IF) buffer (0.1% BSA, 0.2% Triton-X100, 0.1% TWEEN 20 in PBS) and incubated with primary antibodies diluted in IF buffer for 24 to 48 h at room temperature. Organoids were then washed three times with IF buffer for 10 min each with gentle rocking and incubated with secondary antibodies for 24 to 48 h at 4°C in the dark. Following secondary antibody incubation, organoids were washed 3X times for 5 min each with PBS. For imaging, organoids were either transferred to 4-well imaging chambers and imaged in PBS or mounted in VectaShield mounting medium on glass slides, sealed with coverslips, and imaged using Airyscan microscopy. The following primary and secondary antibodies and dilutions were used: Rabbit Phospho-FAK (Invitrogen, #700255; 1:100), mouse anti-α-tubulin (Abcam, #ab7291; 1:200), rat anti-ZO-1 (Stanta Cruz, #33725; 1:200), Phalloidin-Alexa Fluor 568 (Thermo, #A12380; 1:300), DAPI (Thermo, # 62248; 1:500), Alexa Fluor 568 Goat anti-Rat IgG (H+L) (Invitrogen, #A-11077; 1:300), Alexa Fluor 568 Goat anti-Rabbit IgG (H+L) (Invitrogen, # A-11011; 1:300), and Alexa Fluor 568 Goat anti-Mouse IgG (H+L) (Invitrogen, #A-11004; 1:300).

### Tissue sectioning

Small intestines isolated from GFP-M2 KI mice were flushed thoroughly with cold PBS to remove luminal contents, freed of visible connective tissue, fixed overnight in 4% PFA, washed 3X with PBS, and incubated overnight in 30% sucrose in PBS for cryoprotection. Tissue was then cut into appropriate-sized pieces and embedded in OCT compound (Tissue-Tek, #4583). Cryosections 10 to 20 µm thick were prepared using a Leica CM1860 UV cryostat, collected onto charged microscope slides, and imaged.

### Microscopy

AiryScan imaging was performed using Zeiss LSM 880 and 980 confocal microscopes equipped with AirScan, either a 63 X 1.4 NA oil objective or a 20 X air objective, and a temperature control chamber set to 37°C and 5% CO_2_. Airyscan image reconstruction was performed using Zeiss ZEN imaging software. Post-acquisition analysis was performed using Zeiss ZEN software and Fiji. Imaging of fixed GFP-M2A KI organoids stained with Phalloidin and anti-ZO-1 antibody was performed using a Zeiss Light Sheet LLS7 microscope equipped with a 50 X NA 1.31 water immersion objective. Post-acquisition analysis was performed using Fiji and Imaris.

### Blebbistatin treatment

All Blebbistatin experiments were performed using para-nitro-Blebbistatin (pnBB; Cayman Chemical, #24171) at a final concentration of 30 µM with the exception of the time lapse imaging associated with Figure 10, where we used 30 µM para-amino-Blebbistatin (paBB; Cayman Chemical, #22699). For the long term, time lapse Airyscan imaging associated with Figure 7, RFP-LifeAct expressing organoids were treated with pnBB at the start of imaging (0 h) and imaged at 1 h intervals using a 20 X air objective. In addition, the medium was replaced with fresh medium containing pnBB every 36 h to maintain drug exposure. For quantification of orthogonal cell divisions and interkinetic nuclear migration (INM), GFP-H2B expressing organoids and RFP-neuromodulin expressing organoids were treated with pnBB for 24 h and then imaged at 5 min intervals to obtain the data for analysis.

### Laser ablation

For the laser ablation experiments, GFP-M2A KI organoids were seeded into CellVis chambers (CellVis, #C4-1.5H-N) and imaged on a Nikon Eclipse Ti2-E microscope equipped with a 60 X 1.4 NA oil-immersion objective, a CSU-W1 spinning-disk confocal scanner (Yokogawa), an ORCA-Fusion digital CMOS camera (Hamamatsu Photonics), a 488 nm laser line, and a UV ablation setup employing a 355 nm Q-Switched DPSS 40 uJ free space pulsed laser with an electronic attenuator controlled by NIS-Elements imaging software. The incubator chamber enclosing the sample was maintained at 37°C and 5% CO₂. A single ablation was performed per organoid, after which organoids were imaged for at least 10 min at 1-sec intervals to capture cell-shape deformations. Images were analyzed using Fiji/ImageJ. To quantify cell-shape deformation, the cell surface was assumed to be locally spherical, and circular fits were applied independently to the ablated side (radius R1; magenta) and the opposite, non-ablated side (radius R2; blue) (see Figure 5D). Local curvature (κ) was calculated as κ = 1/R.

### Particle Image Velocimetry (PIV)

Laser ablation-induced cellular displacement was quantified using a publicly available Particle Image Velocimetry (PIV) algorithm implemented as a Fiji/ImageJ plugin. PIV analysis was performed on sequential time-lapse images to calculate local displacement vectors between consecutive frames. The PIV interrogation and search window sizes for the first, second, and third passes were set to 95 and 190, 65 and 130, and 35 and 70 pixels, respectively. Correlation, noise, and threshold parameters were set to 0.6, 0.20, and 5, respectively. Images were acquired at 6s intervals, and cumulative displacement over the 80s period following laser ablation was calculated from the sequential frame-to-frame displacement measurements.

### Calculating the axial orientation of myosin bipolar filaments within the metaphase myosin filament cage

The longitudinal axis of GFP-M2A bipolar filaments within the metaphase myosin filament cage (see Figure 4, Panel C), which appear in super resolution images as doublets of GFP spaced ∼300 nm apart, was determined by drawing a line connecting the centers of the two GFP signals. To calculate the axial orientation of the filaments comprising the metaphase myosin filament cage, the angle (θ) between the filament’s longitudinal axis and the dividing cell’s apical–basal axis was measured. The more the filament’s longitudinal axis aligns with the dividing cell’s apical–basal axis, the smaller θ becomes. Within a single focal plane of an AiryScan image, ten cortical strings consisting of aligned bipolar filaments (approximately 8 μm in lateral width) were selected per cell, and a total of 10 metaphase cells were scored.

### Quantification of M2A filament orientation fields in metaphase cells

The orientations of fields of GFP-M2A filaments in metaphase cell cortices (see Figure 4, Panels D-F) were measured using the public-domain program OrientationJ implemented as a Fiji/ImageJ plugin (Rezakhaniha et al., *Biomech Model Mechanobiol*, 2012)[81]. OrientationJ computes orientation, energy, and coherency maps using structure-tensor analysis and generates weighted orientation histograms. Maximum-intensity-projected Airyscan images of metaphase cells were used for orientation analysis. The structure tensor was computed using a Gaussian window of 2 pixels and a cubic-spline gradient. Orientation distributions presented as frequency histograms binned in 10° increments and ranging from −90° to +90° were calculated relative to either the cell’s apical/basal axis for cells dividing within the epithelium or the horizontal axis of the image for cells dividing in isolation. For comparison across conditions, corresponding orientation bins were summed and normalized to the total frequency.

### Statistical analysis

Data are given as mean ± SD. The statistical significance of the difference between two sets of data was evaluated using the two-tailed, unpaired Student’s t-test. Statistical significance was accepted at P < 0.05 and indicated as follows: * = *P* < 0.05, ** = *P* < 0.01, *** = *P* < 0.001.

## ACKOWLEDGMENTS

The authors thank the NHLBI Light Microscopy Core for providing access to the Zeiss 980 and LLS7 microscopes, the NIAMS Light Imaging Core for providing access to the Leica cryostat, Dr. Takashi Akera for providing access to his laser ablation setup, and the Animal Surgery and Resources Core for providing access to facilities and technical support.

## AUTHOR CONTRIBUTIONS

**S.Y.** and **J.A.H.** conceptualized the project, developed the overall research strategy, interpreted the results, and collaborated on drafting the manuscript. **J.A.H.** provided scientific guidance, supervision, and mentorship throughout the project. **S.Y.** led the project, designed and performed the majority of experiments, developed quantitative analysis approaches, provided mechanistic interpretations of the experimental findings, and prepared the figures. **K.R.** established the organoid system in the lab, performed the initial M2A knockdown experiments, and was the first to see the M2A cage-like structure. **A.G**. and **Y.Y.** contributed to the design and preparation of lentiviral vectors and plasmids. **G.P**., **L.B.**, **C.A.**, and **A.P.** assisted with intestinal organoid culture, maintenance, and optimization of various organoid-related procedures. **D.S.** provided the GFP-M2A KI mice. **X.W.** contributed expertise in AiryScan and Lattice Light Sheet microscopy, image analyses, and data interpretation. All of the authors reviewed and approved the final manuscript.

## DISCLOSURES

The authors declare no competing interests.

*Disclaimer: This research was supported by the Intramural Research Program of the National Institutes of Health (NIH). The contributions of the NIH author(s) are considered Works of the United States Government. The findings and conclusions presented in this paper are those of the author(s) and do not necessarily reflect the views of the NIH or the U.S. Department of Health and Human Services*.

## Notes

### Competing Interest Statement

The authors have declared no competing interest.

