## Supplementry Figures for "A myosin filament cage drives multiple aspects of mitosis unique to cells dividing within a polarized epithelium"

__List of Supplementary Figures__

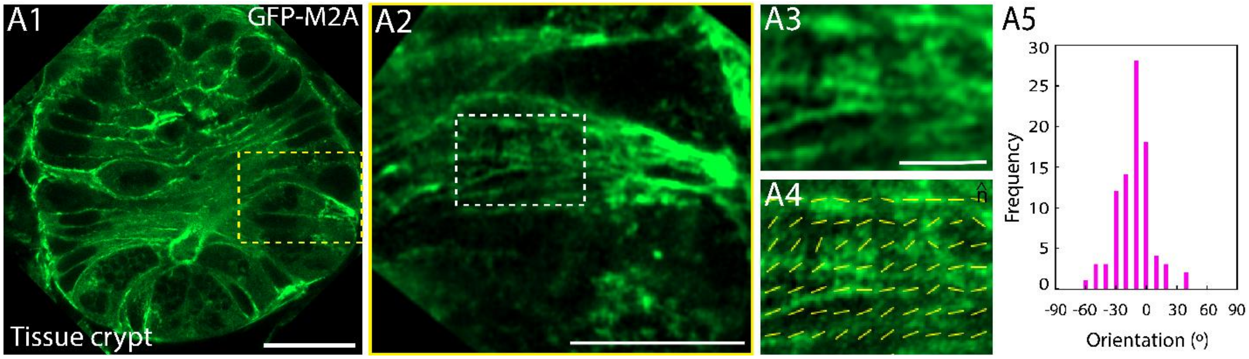


**Figure S1: Orientations of the M2A filaments in the cortex of metaphase cells present within the intestinal tissue crypt.** (A1) Representative, single-plane, Airyscan image of an intestinal crypt within fixed tissue from the GFP-M2A KI mouse (the crypt is sectioned crosswise). The yellow box indicates a mitotic cell. (A2) A maximum intensity projection of the dividing cell boxed in A1 (oriented such that the cell's apical/basal axis is horizontal). (A3) An enlargement of the region marked with a white box in A2. (A4) The long axis of all the M2A filaments within the boxed region in A3 are marked with yellow lines. (A5) Histogram of the orientations of all the marked filaments in D3 relative to the dividing cell's apical/basal axis. An orientation of 0° is parallel to this axis, while values of -90° and +90° are perpendicular and in opposite orientations relative to the cell's apical/basal axis. Scale bars: (A1) 20 µm, (A2) 10 µm, (A3) 5 µm.

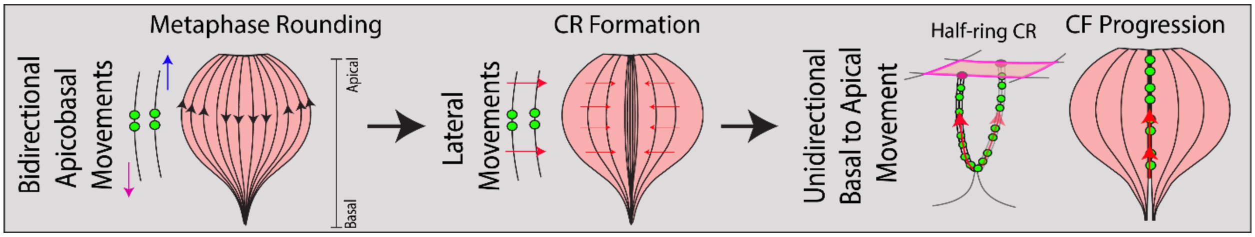


**Figure S2: Schematic depicting the coordinated movements of the M2A filament strings comprising the metaphase M2A filament cage during cell division within a polarized epithelium.** The M2A filament strings comprising the metaphase cage undergo three concerted movements during mitosis: bidirectional apicobasal movements during metaphase cell rounding, lateral coalescence during contractile ring (CR) formation, and unidirectional, basal-to-apical motions during cleavage furrow (CF) progression in the form of a contractile "half ring".

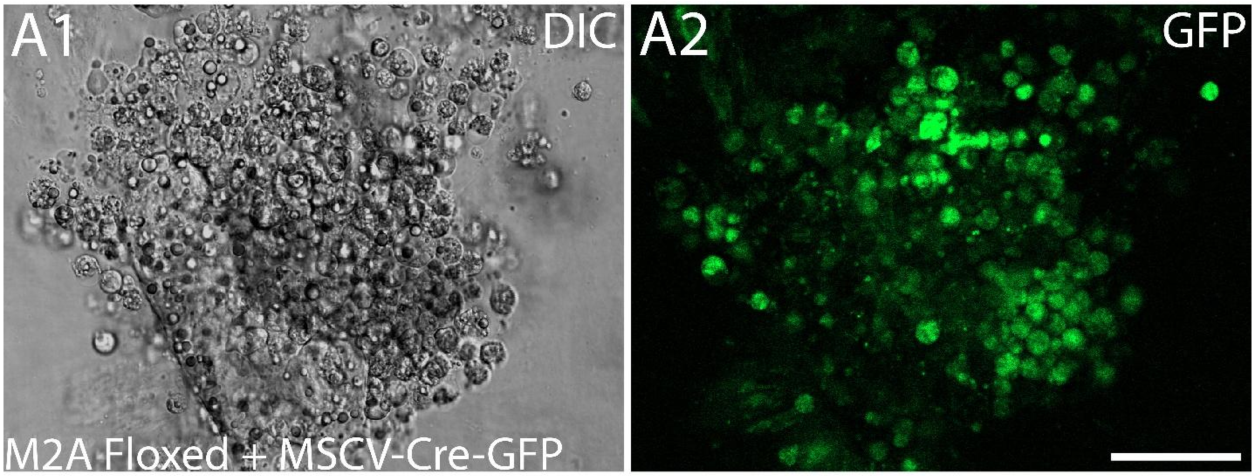


**Figure S3. Complete loss of M2A compromises intestinal organoid viability and epithelial integrity.** Shown is a transmitted light image (A1) of an organoid made from the floxed M2A cKO mouse that was treated with a MSCV-Cre-GFP retrovirus to excise the M2A heavy chain gene. (A2) The corresponding green signal indicating expression of the Cre recombinase. Scale bar, 50 µm.

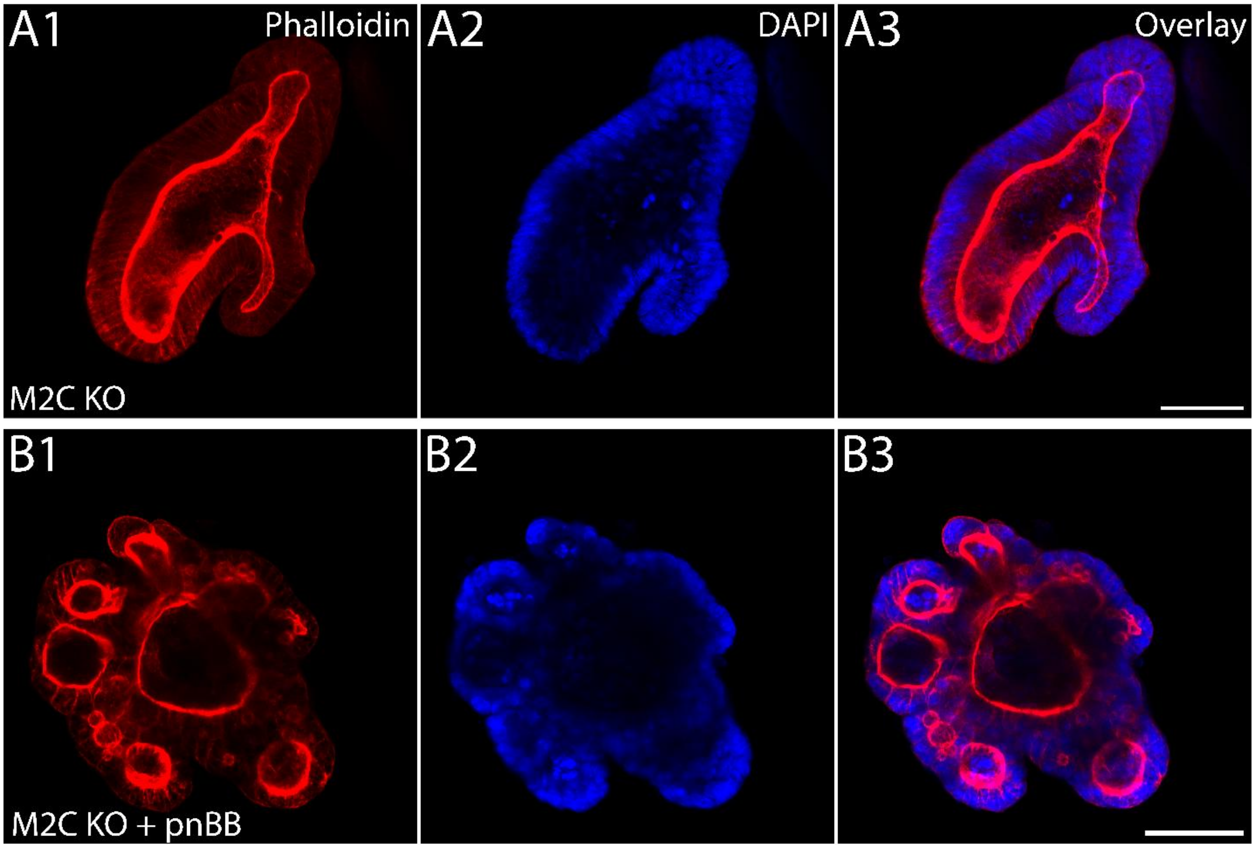


**Figure S4: M2C KO organoids possess a single lumen unless treated with pnBB to inhibit M2A.** (A1-A3) A maximum intensity projected Airyscan image of a representative M2C KO organoid stained with Phalloidin (A1) and DAPI (A2) (the overlaid image is shown in A3) showing a single lumen. (B1-B3) As in A1-A3 except that this M2C KO organoid was treated for 48 hrs with pnBB, which results in the multi-lumen phenotype. Scale bars, 50 µm.
